# Deuterium labeling enables proteome wide turnover kinetics analysis in cell culture

**DOI:** 10.1101/2025.01.30.635596

**Authors:** Lorena Alamillo, Dominic C. M. Ng, Jordan Currie, Alexander Black, Boomathi Pandi, Vyshnavi Manda, Jay Pavelka, Peyton Schaal, Joshua G. Travers, Timothy A. McKinsey, Maggie P. Y. Lam, Edward Lau

**Affiliations:** Department of Medicine; Consortium for Fibrosis Research & Translation; Department of Biochemistry and Molecular Genetics University of Colorado School of Medicine Aurora, CO 80045, USA

**Author notes:** Equal Contributions (Lorena Alamillo, Dominic C. M. Ng, Jordan Currie). **Lead Contact**: Edward Lau, Ph.D.

**Keywords:** Protein turnover, turnover kinetics, heavy water, hiPSC, protein synthesis, degron, secretome, mass spectrometry

## Abstract

The half-life of proteins is tightly regulated and underlies many cellular processes. It remains unclear the extent to which proteins are dynamically synthesized and degraded in different cell types and cell states. We introduce an improved D_2_O labeling workflow and apply it to examine the landscape of protein turnover in pluripotent and differentiating human induced pluripotent stem cells (hiPSC). The majority of hiPSC proteins show minimal turnover beyond cell doubling rates, but we also identify over 100 new fast-turnover proteins not previously described as short-lived. These include proteins that function in cell division and cell cycle checkpoints, that are enriched in APC/C and SPOP degrons, and that are depleted upon pluripotency exit. Differentiation rapidly shifts the set of fast-turnover proteins toward including RNA binding and splicing proteins. The ability to identify fast-turnover proteins in different cell cultures also facilitates secretome analysis, as exemplified by studies of hiPSC-derived cardiac myocytes and primary human cardiac fibroblasts. The presented workflow is broadly applicable to protein turnover studies in diverse primary, pluripotent, and transformed cells.

## Introduction

Protein abundance in cells is governed by a balance between synthesis and degradation. The half-life of a protein pool regulates its homeostasis and function, influencing diverse biological processes from cell cycle to gene regulation and stress response ^1–5^. Unlike homeostatic tissues in vivo, rapidly proliferating cells in culture can remove proteins through two principal mechanisms: passive dilution to daughter cells, as well as active degradation by proteolysis ^6,7^. Proteins regulated by the latter undergo excess synthesis and degradation cycles beyond what is required for cell doubling. Prior efforts have mapped hundreds of short-lived or rapidly degraded proteins with half-life < 8 hours to identify the regulatory roles of protein degradation and find therapeutic targets for proteolytic interventions ^8,9^. In parallel, continued protein translation has been shown to be required to maintain open chromatin, a hallmark of pluripotent cells ^10^. The extent to which the identity, timing, and kinetics of these fast-turnover proteins differ across cell types remain unclear. Identifying the proteins that undergo targeted synthesis and degradation in pluripotent cells may shed light on intervention targets relevant to cell proliferation.

Protein turnover measurements in cell culture have largely relied on cycloheximide inhibition of protein synthesis ^8,9^, or dynamic stable isotope labeling by amino acid in cell culture (SILAC)^11–13^. Heavy water (D_2_O) labeling mass spectrometry provides an alternative to isotope-labeled amino acids for tracing protein turnover, and has the potential to offer compatibility with standard cell culture conditions, eliminating possible secondary effects of cycloheximide or the need for specialized SILAC medium formulations. Upon D_2_O labeling, deuterium is rapidly incorporated at stable C–H bonds in non-essential amino acids during their biosynthesis and metabolism ^14–16^. Labeled amino acids are subsequently incorporated into nascent protein chains to enable turnover quantification. D_2_O labeling has been successfully applied to trace protein synthesis in rodents ^14–19^, non-human primates ^14^, and humans ^16,20–22^; but its adoption to cell culture remains under-realized. A key variable in analyzing D_2_O labeling data to derive kinetics information is the number of deuterium accessible sites in each identified peptide, which can be calculated in animal experiments but has not been deeply investigated in cultured cells. Although D_2_O in culture media equilibrates with hydrogen atoms in cellular amino acids ^23,24^, prior work has largely assumed effective protein labeling only at Gly and Ala residues, despite data discrepancies ^25^. It therefore remains unclear to what extent D_2_O labels various cellular proteins, especially given the high concentration of free amino acids in culture media ^24,25^. This barrier complicates the interpretation of labeled spectra to extract kinetics information and hinders applications. Hence, few studies have applied D_2_O to measure individual protein turnover in cells, and their scope was confined to a handful of pre-selected peptides rather than proteome-scale investigations ^17,25,26^.

In this study, we developed an updated workflow using D_2_O labeling to measure proteome-wide protein turnover rates in cell culture. To enable interpretation of labeled mass spectrometry data, we determined the extent of deuterium enrichment in all 20 proteinogenic amino acids following D_2_O enrichment, by using machine learning strategies and experimental calibration with cells of known labeling proportions. The established per-amino acid enrichment values show clear differences from those used in animal studies or commonly assumed in earlier cell culture experiments, and enable precise turnover calculations for virtually any protein sequence. We demonstrate the versatility and robustness of deuterium labeling in two separate applications, first to map the protein turnover landscape of human induced pluripotent stem cells (hiPSC) under maintained pluripotency and directed differentiation, and secondly to characterize the secretome of human cardiac cells under baseline or stressed conditions. The results demonstrate D_2_O labeling is a flexible and economic method to broadly assess protein kinetics in diverse systems.

## Design

Dynamic SILAC coupled to mass spectrometry is used to measure protein turnover dynamics in cell culture, but requires altering culture medium composition and may not label some peptides. Building on prior work by us and others to develop software and analytical methods for turnover studies, we describe a simple and convenient alternative to mix culture medium with 6% D_2_O (heavy water) for analyzing protein turnover kinetics and secretome flux at proteome-scale. D_2_O labels primarily non-essential amino acids not supplemented in media to enable mass spectrometry analysis of isotope incorporation rates. Addressing a critical gap, we determined the deuterium-accessible atoms on all 20 proteinogenic amino acids across multiple cell lines.

This allows accurate interpretation of D_2_O labeled mass spectra to extract protein kinetics information, and enables proteome-scale turnover measurements from single or multi time-point continuous labeling designs.

## Results

### D_2_O incorporates deuterium into multiple TCA cycle amino acids in cell culture

D_2_O labeling leads to a gradual shift in the isotope profiles of peptides, which is a function of the excess deuterium enrichment (*p*), the number of deuterium-accessible labeling sites in a peptide (*S*_pep_), and the fraction of newly synthesized peptide (θ) in the sample (**Figure 1A**). Finding the parameter of interest (θ) needed to calculate protein half-life therefore requires knowing the number of deuterium-accessible atoms in each peptide. In animal experiments, the conventional method uses the sum of per-amino acid labeling sites in the sequence, with commonly used values measured from mouse tritium labeling ^27^.

**Figure 1:**
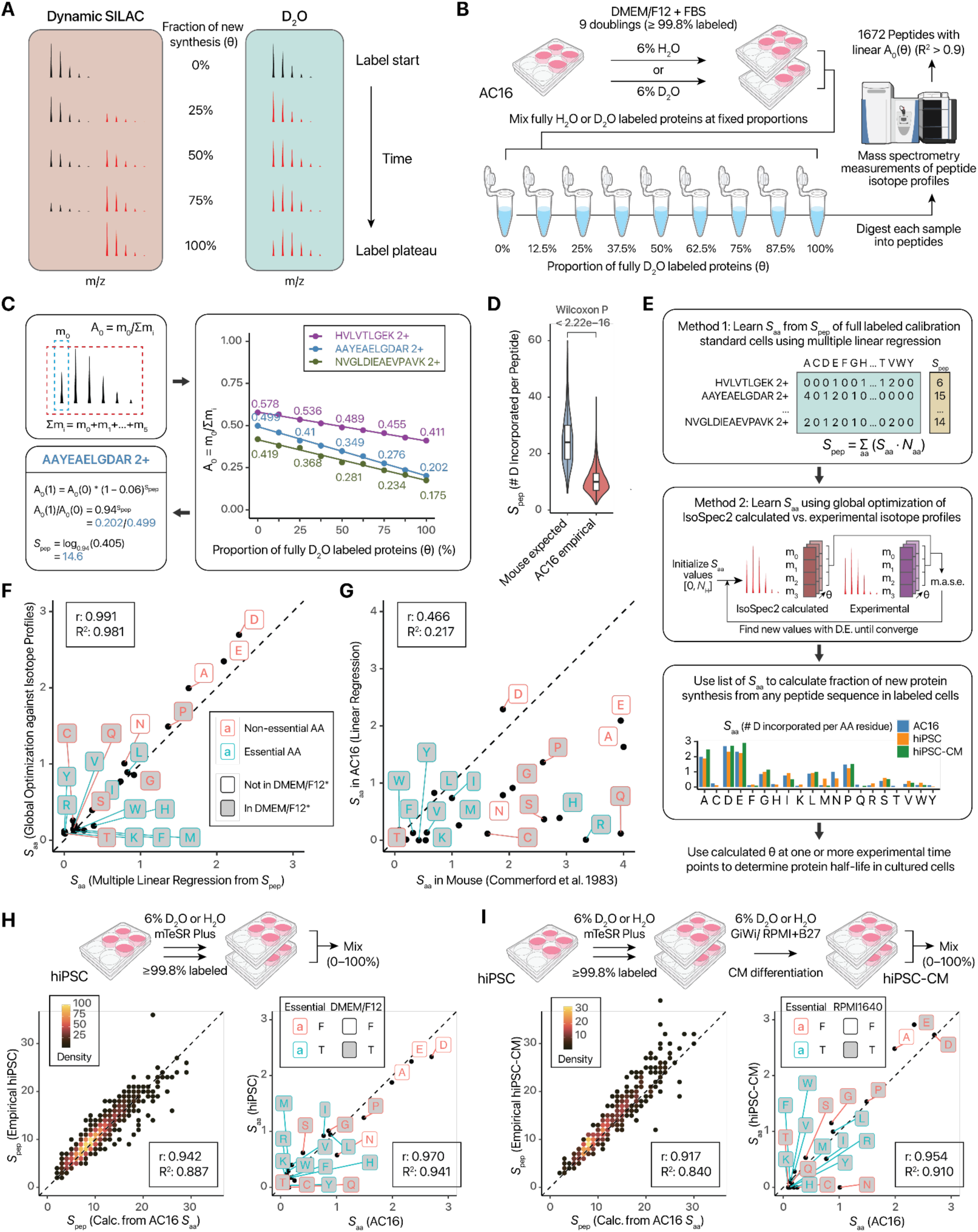
D_2_O labeling incorporates deuterium into proteins in cell culture with residue specificity. **A.** Mass shifts of peptide isotope envelopes following SILAC and D_2_O labeling **B.** Calibration standard cells are cultured in media with 6% H_2_O or D_2_O for 9 doublings to yield unlabeled and fully labeled proteins, which are then mixed in fixed proportion and analyzed by LC-MS/MS. **C.** Calculation of peptide labeling sites (*S*_pep_) from standards. The proportional abundance of the m_0_ peak, which contains no heavy isotopes in any atom center, decreases as the proportion of labeled protein scales from 0–100%. Right: line plots of three peptides measured in our experiments. The position of A(0) is a function of peptide mass/length whereas the decrease depends on numbers of D accessible labeling sites. **D.** Violin/boxplot of peptide labeling sites calculated from the m_0_/m_i_ ratio of the labeling standard samples, highlighting the difference (Wilcoxon P <2.2e–16) to numbers calculated from labeling sites used in adult mouse experiments. **E.** Prediction of labeling sites per amino acid (*S*_aa_). Method 1 uses multiple linear regression to learn *S*_aa_ from *S*_pep_. Method 2 uses global optimization based on differential evolution (D.E.) to find the *S*_aa_ values that lead to IsoSpec2 isotopomer profiles resembling experimental data, minimizing median absolute scaled errors (m.a.s.e.). Barcharts show the number of label-accessible hydrogens per amino acid across three cell types. **F.** Scatterplot showing the high correlation of *S*_aa_ values (*r*: 0.991) predicted using multiple linear regression against empirical *S*_pep_, and using global optimization against isotopomer profiles. Dashed line: 1:1. *: Label shading denotes whether amino acids are supplemented in DMEM/F12 at appreciable concentration (tryptophan at any concentration, plus other amino acids at > 10 mg/L). DMEM/F12 additionally contains low levels (< 10 mg/L) of alanine, aspartic acids, asparagine, glutamic acids from the F12 nutrient mix that are not marked on the graph. **G.** Scatterplot comparing learned *S*_aa_ values in AC16 against in vivo mouse ^3^H labeling sites from Commerford et al. **H.** Labeling sites in hiPSC cultured in MTeSR Plus. Left: AC16-derived *S*_aa_ values accurately predict measured *S*_pep_ in hiPSC. Right: highly comparable predicted *S*_aa_ (*r*: 0.97) between two cell types. Label shading denotes whether amino acids are supplemented in DMEM/F12 in lieu of mTeSR Plus. **I.** As in H, but for hiPSC-CM. hiPSC was passaged in 6% H_2_O or D_2_O prior to differentiation into hiPSC-CM. Label shading denotes whether amino acids are supplemented in RPMI-1640 media (at ≥ 10 mg/L with the exception of L-tryptophan)

To acquire these values in cultured human cells, we take advantage of a complete labeling calibration standard that we recently generated ^28^, by culturing human AC16 cells in DMEM-F12 basal media diluted with 6% D_2_O, supplemented with 10% FBS, for at least 9 doublings (**Figure 1B**). This guarantees ≥99.8% complete labeling from cell division even in the absence of additional protein degradation, i.e., all proteins are fully labeled with 6% deuterium. This fully labeled pool is then mixed with unlabeled cell lysate at fixed proportions (0%, 12.5%, 25%, 37.5%, 50%, 62.5%, 75%, 87.5%, 100%) to establish the ground-truth fractional synthesis in each sample (**Figure 1B**). From the fully labeled samples, per-peptide labeling sites can be calculated by considering the proportion of peptide molecules that remain unlabeled, even without knowing the individual amino acid labeling sites (Methods; **Figure 1C**). As expected, D_2_O incorporates deuterium into peptide sequences in cell culture, leading to a drop of the ratio of the monoisotopic peak over the entire isotope envelope (*A*_0_); however, considerably fewer atoms are incorporated than in animals (i.e., lower drop in *A*_0_ from 0–100% labeling), with median 9.7 sites per peptide, vs. 24.6 when estimated using adult mouse (in vivo) labeling sites (**Figure 1D**). Training a multiple linear regression model, we then learned the number of deuterium-accessible sites from the total labeling sites of each peptide and their amino acid composition (**Figure 1E; Table S1**). In parallel, we corroborate the labeling sites with a complementary machine learning strategy that does not rely on the fitted *A*_0_ information, by performing nonlinear global optimization to minimize the differences between the empirical isotope profiles and the predicted profiles calculated from a fine structure calculator IsoSpec2 ^29^ (Methods; **Figure 1E**). The two strategies returned near-identical values (r: 0.991; R^2^: 0.981), indicating we are able to acquire internally consistent information on the extent of deuterium labeling in these cells (**Figure 1F**). The results highlight that amino acid residues Asp, Glu, Ala, Pro, Gly, Leu, Ile, Asn are relatively accessible to D_2_O labeling, whereas compared to the adult animals, Gln, Arg, Ser, His are inaccessible to D_2_O labeling in cell culture (**Figure 1G**, **Figure 1E**). Therefore, D_2_O is able to exchange with hydrogen in multiple TCA-cycle derived amino acids; notably Asp, Ala, Pro, Glu. At the same time, the models predict ∼2 exchangeable hydrogens in alanine, which is considerably lower than the 4 in vivo or previously assumed in cell culture (**Table S1**).

We next extended the investigation to generating new fully-labeled calibration standards in two additional cell types, namely human induced pluripotent stem cells (hiPSC), and hiPSC-derived cardiomyocytes (hiPSC-CM). Both are widely employed as physiologically-relevant cell models for drug discovery and disease mechanism studies. Notably, hiPSC are commonly cultured in specialized media (e.g., mTeSR Plus) for which SILAC-compatible depleted basal media is not commercially available, therefore necessitating custom media for dynamic SILAC and providing a compelling use case for the convenience and flexibility of D_2_O labeling as an alternative method. To generate the calibration cells, hiPSC were grown to ≥10 doublings in mTeSR Plus. To procure fully-labeled non-proliferative hiPSC-CM, hiPSC were fully labeled in D_2_O prior to CM differentiation. The empirical peptide labeling sites in hiPSC are similar to what would be predicted using the AC16-derived cell culture values (**Figure 1H)** (Pearson’s correlation r: 0.942, R^2^: 0.887) and similarly for hiPSC-CM (**Figure 1I**) (r: 0.917, R^2^: 0.840). The trained linear regression models likewise predicted similar numbers of labeling sites (r: 0.970 for hiPSC, r: 0.954 for hiPSC-CM) (**Figure 1H**, **Figure 1I**). Notably different is the labeling of Asn, which incorporates deuterium in AC16 under DMEM/F12 with FBS, but to a lesser degree in hiPSC under MTeSR and absent in hiPSC-CM under RPMI-1640 with B27, which contains supplemented Asn. On the other hand, multiple non-essential TCA cycle amino acids remain effectively labeled by D_2_O in hiPSC-CM, despite RPMI-1640 being a complete medium that contains Asp, Glu, and Pro, indicating the primary source of these amino acids for protein production likely lies in endogenous biosynthesis rather than external uptake. Taken together, the predicted labeling sites show a high degree of consistency, while reflecting potential differences in cell types or culture media compositions.

### Amino acid analysis corroborates the determined D_2_O labeling sites

To further verify per-residue D_2_O labeling, we performed acid hydrolysis to experimentally release free amino acids from proteins in unlabeled and fully labeled AC16 cells, then measure the deuterium-tagged amino acids directly (**Figure 2A**). Direct infusion high-resolution MS allows the mass defect differences between ^13^C_1_-containing and D_1_-containing amino acids to be distinguished, and their respective relative abundance to the monoisotopic peaks to be compared to the calculated isotope profiles. The relative intensity of the signals corresponding to D_1_-alanine (m/z 91.0621, +1 charge) is consistent with predictions in cell culture (1–2 labeling sites, vs. 4 in adult mice in vivo) (**Figure 2B–C**). The measurements are not biased by the transient time of the Orbitrap mass spectrometer, as similar results are obtained at different mass resolutions (**Figure S1**). On the other hand, the absence of signals corresponding to D_1_-arginine (m/z 176.1252, +1 charge) is consistent with its predicted absence of deuterium-accessible labeling sites in cell culture (**Figure 2D–E**); whereas amino acid analysis also supports 1–2 labeling sites on average in proline, and <1 site in glycine and leucine/isoleucine (**Table S2; Figure 2F**). Taken together, data in three cell types (transformed AC16, pluripotent hiPSC, terminally differentiated hiPSC-CM) highlight that deuterium incorporation sites can be reliably determined, supporting the use of D_2_O labeling for protein turnover studies in cell culture with different cell types and media. The results here provide the values of labeling sites that are consistent for experiments in different cell types, and a general strategy to find cell type-specific labeling sites.

**Figure 2.**
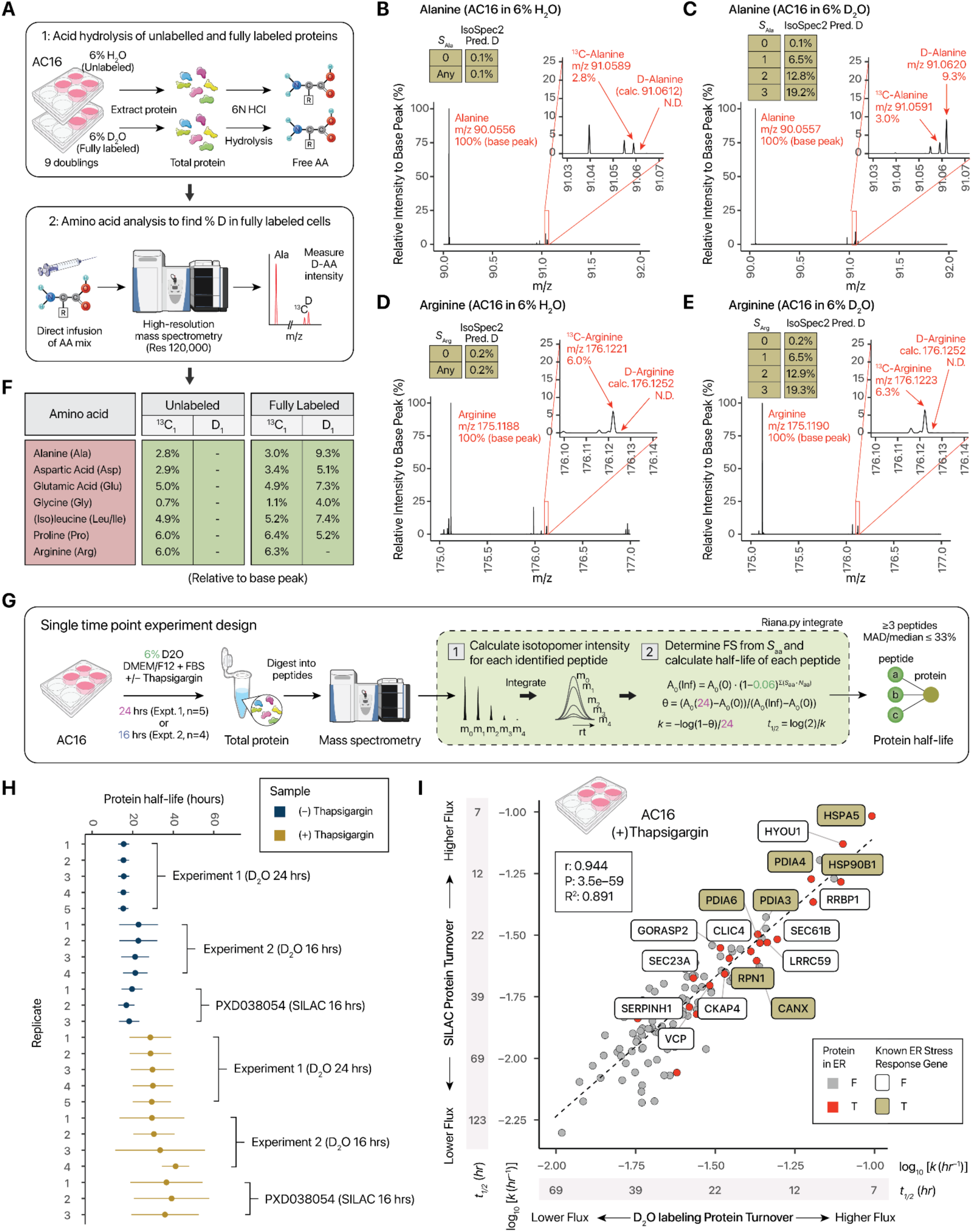
Experimental validation of deuterium incorporation sites and single time-point kinetics analysis. **A.** Mass spectrometry experimental validation of predicted labeling sites in cell culture. At 120,000 resolution, ^13^C-amino acid and D-amino acid peaks can be resolved by their mass differences from mass defects, allowing their relative intensities to be compared with expectation under different D-accessible labeling site predictions. **B.** Mass spectra from acid hydrolysis of unlabeled samples, showing no detectable D-alanine. N.D.: not detected **C.** As in B, but for alanine in fully labeled samples, showing detectable D-alanine at m/z 91.0612 with a relative intensity of 9.3% to monoisotopic (m/z 90.0550) alanine. The Inset table shows IsoSpec2 predicted D-alanine relative intensity (0.1%, 6.5%, 12.8%, and 19.2%) relative to the monoisotopic peak if *S*_Ala_ were 0, 1, 2, and 3, respectively. The data is consistent with 1 – 2 labeling sites in cell culture as opposed to 4 labeling sites in adult mice in vivo. **D.** As in B, but for Arg. **E.** As in C, but for Arg, showing no detectable D-Arg (m/z 176.1252) in cell culture. **F.** Table of D-AA proportion in Ala, Asp, Glu, Gly, Leu/Ile, Pro, and Arg in unlabeled and fully labeled samples, as measured by amino acid analysis using high-resolution MS. Asn and Gln are deamidated during acid hydrolysis _33_, hence the measured Asp and Glu values are a function of both Asp/Asn and Glu/Gln. **G.** Application of D_2_O labeling to a cell culture model of ER stress. proteome-wide protein kinetics in AC16 cells. **H.** Pointranges of protein half-life measured by D_2_O and SILAC in AC16, showing comparable ranges between methods. D_2_O recapitulated a slowdown in cell proliferation and protein synthesis upon thapsigargin-induced ER stress. Error bars: s.d. **I.** Scatter plot between D_2_O and SILAC measurements upon 1 µM thapsigargin treatment, showing strong correlation (r: 0.94) and revealing stress response genes with high turnover flux upon ER stress.

### D_2_O labeling enables single-point measurement of protein synthesis kinetics in cells

We next evaluated whether the labeling sites support interpretation of isotopomer envelopes to reflect accurate turnover rates on a proteome-scale. To do so, we first applied dynamic D_2_O labeling to proliferating human AC16 cells for a fixed period of time, then quantified isotope incorporation from a single time point (**Figure 2G**). Because the determined D_2_O labeling sites enable the asymptote of mass shifts to be calculated, the label incorporation kinetic curve of a single protein can be completely defined by a single variable, namely the apparent turnover rate of the protein. This therefore allows rise-to-plateau kinetics to be analyzed from as few as one time point, provided the sampling time point is within the informative region of the curve (i.e., an appreciate amount of isotope has been incorporated, and the curve has not plateaued) ^30,31^.

Although single-point sampling approaches are challenged by the high variability of derived turnover rates – due to the lack of filtering from goodness-of-fit and the limited dynamic range that can be sampled – it can be useful for evaluating whether the isotope envelope is quantified accurately to reflect expected median half-life of proteins. It also has important applications for when multi-time point collection is infeasible, such as in single-cell measurements.

Under baseline conditions, AC16 cells proliferate quickly with a doubling time of ∼16–20 hours, and protein isotope incorporation is expected to be driven by the doubling time under the particular culture condition. Applying 6% D_2_O to AC16 cells for 24 hours (n=5) then quantifying individual protein turnover rates, we measured a proteome-wide median half-life of 16.7–17.5 hours (**Figure 2H; Table S3**), in line with prior data (∼16–20 hours) ^32^. We did not observe a change in proliferation rate upon diluting the media with low-dose (6%) D_2_O, which is consistent with prior reports on the minimal effect on cell growth ^25^. For comparison, we applied the single-point sampling approach to non-proliferative hiPSC-CM upon 48 hours of dynamic labeling. The experiment yielded the turnover rates of 1406 proteins with at least 3 measured peptides and intraprotein c.v. ≤ 33%. The hiPSC-CM data shows a distribution of slower median turnover rates that roughly follows a log-normal distribution, with a median half-life of 53.9 hours (**Table S4**; **Figure S2**). In parallel, we applied D_2_O labeling to primary human cardiac fibroblasts (hCF), obtained from non-failing donor hearts not suitable for transplant, and labeled with D_2_O for 48 hours. Because primary cells are replicatively limited, they can only be cultured for a finite number of passages prior to senescence. We therefore did not pursue a fully labeled calibration standard, and instead opted to apply the deuterium accessible site values from hiPSC-CM toward the analysis. (**Table S5**). The hCF proteins show a median half-life of 55.1 hours, follow a log-normal distribution, and show a moderate but significant correlation with hiPSC-CM half-life (r: 0.32, P: 6.3e–10; **Figure S2**). Therefore, the labeling sites support the calculation of a range of isotope incorporation kinetics across cell types.

To evaluate the accuracy of the derived turnover rates, we applied D_2_O labeling to AC16 under ER stress, and compared to SILAC data. Thapsigargin-induced ER stress is known to halt cell proliferation, and we observed a near-twofold reduction in protein half-life upon a single concomitant dose of thapsigargin with D_2_O, to 31.5–34.8 hours. A second set of replicate experiments (n=4), performed by a different experimentalist using a different AC16 cell stock labeled for 16 hours, likewise recapitulates a near-twofold decrease in turnover rates (**Figure 2H**). Whereas overall protein turnover decreases, the span of measured turnover rates also broadens as ER stress response pathways blunt protein synthesis except for some stress response proteins ^32^, allowing correlative comparison with SILAC measurements previously acquired at 16 hours ^32^. The D_2_O labeling data showed a highly robust correlation with dynamic SILAC (r: 0.94; P: 3.5e–59). At the same time, D_2_O labeling successfully captured the accelerated turnover of ER stress response proteins including heat shock protein family A (Hsp70) family 5 (HSPA5/GRP78/BiP), hypoxia up-regulated 1 (HYOU1), and heat shock protein 90 beta family member 1 (HSP90B1/GRP94) (**Figure 2I**; **Table S5**). These results support the applicability of D_2_O labeling in cell culture for measuring proteome-wide protein half-life and interrogating cellular physiological responses from as few as one time point.

### Mapping the proteome-wide turnover kinetics in hiPSC with D_2_O labeling

We next applied D_2_O labeling to a larger-scale time-course experiment, where hiPSC are labeled in 6% D_2_O in MTeSR Plus at 9 biologically independent time points and moreover in 2 replicate time series (**Figure 3A**). Compared to single-point designs, time-course experiments enable a greater dynamic range of turnover rates to be measured, including fast turnover proteins that may have plateaued prior to sampling time. Moreover, the use of non-linear least square curve-fitting allows data filtering based on goodness-of-fit thresholds to remove problematic peptides that do not conform to the kinetic models (e.g., from low abundance peptides). Upon D_2_O labeling, hiPSC proteins show gradual increases in calculated fractional synthesis as existing proteins are replaced by newly synthesized proteins over 24 hours (**Figure 3B**). The samples were processed to quantify 42,865 distinct peptide sequences that are uniquely mappable to 6905 proteins (median 3599 proteins and 17,545 peptides per sample).

**Figure 3.**
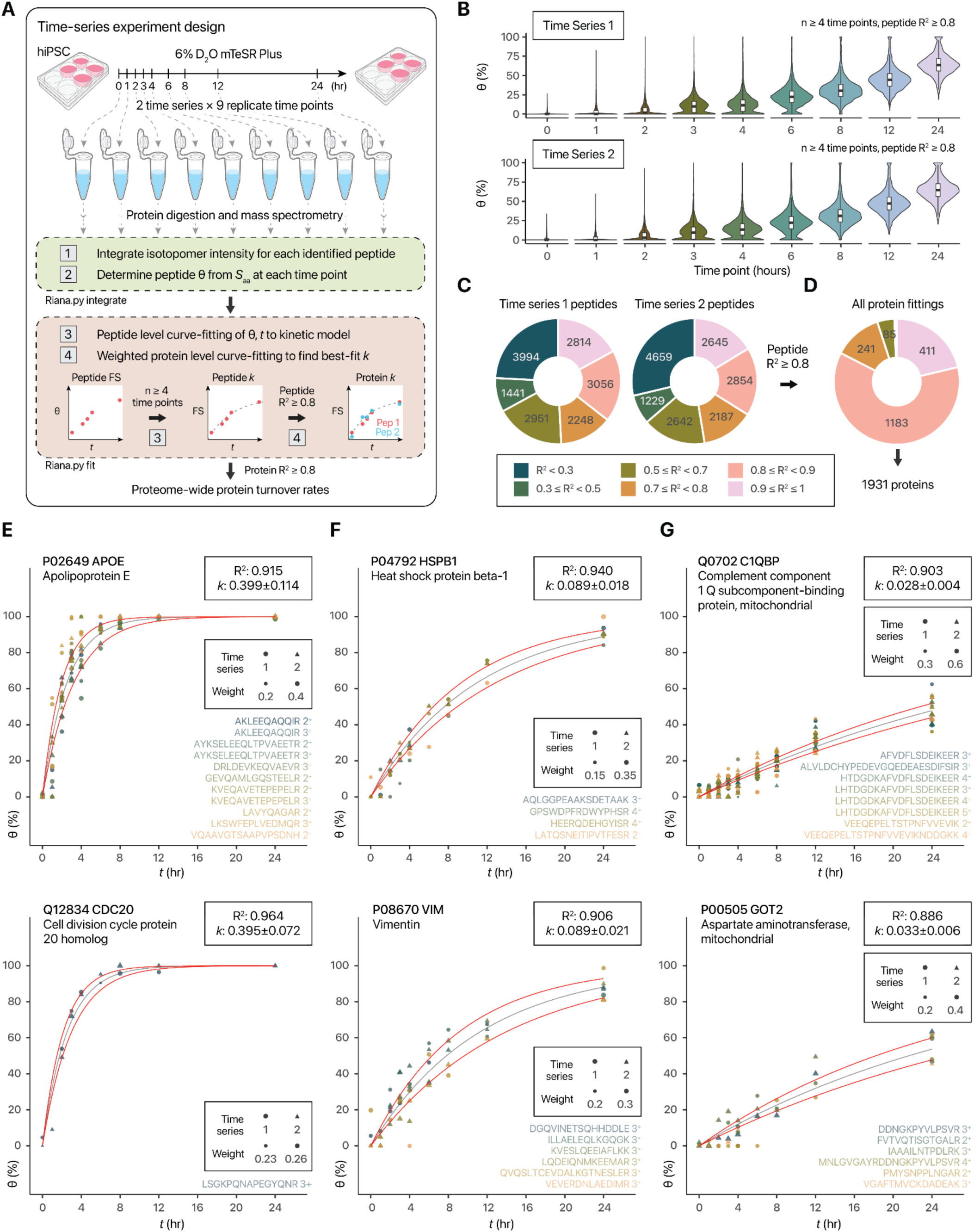
Application of D_2_O labeling to measuring proteome-wide turnover kinetics in hiPSC. **A.** Measurement of protein turnover in hiPSC-CM using D_2_O labeling in a multi-point time-course design. **B.** Rise in fractional synthesis over the course of labeling in two replicate time series of 9 time points each. **C.** Proportion of quantified peptides that fit to a standard kinetic model at different R2 cutoffs. **D.** As in C, but for a second-pass weighted protein-level fitting, after selecting peptides with R^2^ ≥ 0.8 filter. **E.** Isotope incorporation kinetics of three proteins selected to represent the dynamic range of turnover, highlighting two fast-turnover proteins APOE (top) and CDC20 (bottom). Black line: kinetic curve governed by best-fit *k*; red: standard error. **F.** Same as E, but for proteins with moderate turnover HSPB1 (top) and VIM (bottom). **G.** Same as E, but for proteins with low turnover C1QBP (top) and GOT2 (bottom).

We then performed nonlinear optimization of data for each unique peptide ion from all time points to a single exponential kinetic model to derive the best-fit isotope incorporation rate constant *k* while assessing the goodness-of-fit between model and data (**Figure 3A**). This led to the quantification of turnover from 18392 peptides across 2995 proteins. The majority of peptide time series showed good fit to the kinetic model, with about half showing R^2^ ≥ 0.7 and over one-third with R^2^ ≥ 0.8 (**Figure 3C–D**). We used a conservative filter to include only peptides with R^2^ ≥ 0.8 at peptide-level fitting, then combined all peptide data within a protein for a second-pass weighted protein-level fitting to derive protein-level *k*. In total, this led to a higher confidence list of quantified turnover rates of 1931 proteins in hiPSC (**Table S7**). Nearly all proteins retained high R^2^ values, which indicates that the constituent peptides of each protein produce consistent kinetics values, and further strongly corroborates that the deuterium labeling sites allow confident interpretation of spectra to derive fractional synthesis at each time point (**Figure 3E**). The quality of the data is also reflected by a good agreement in protein *k* values when the two time-series are fitted separately (r: 0.76); as well as a median d*k*/*k* (fitting error relative to *k* value) of 24.9%. Thirdly, peptides from the same protein give highly consistent k values, with a median intra-protein peptide geometric coefficient of variance of 20.7%, in line with prior experiments and indicating D2O labeling has similar precision to in vivo D_2_O labeling, as well as SILAC labeling data. The data covers a broad dynamic range of kinetics from fast to slow turnover proteins. Examples of very fast turnover proteins (e.g., *k* >0.1 *hr*^−1^) include APOE and CDC20 (**Figure 3E**); moderate turnover proteins (e.g., *k*: 0.05 – 0.1 *hr*^−1^) include HSPB1 and VIM (**Figure 3F**); low turnover proteins (e.g., *k* < 0.05 *hr*^−1^) include C1QBP and GOT2 (**Figure 3G**). We note that these reported turnover rates (*k*) represent apparent rates of isotope incorporation and do not distinguish the contributions from cell division rates, although this can be readily corrected in the kinetic model by subtracting the doubling rate from the rate constant.

Comparing the turnover rate differences with intracellular structure and functional categories, we find that mitochondrial proteins have on average longer half-life, with a majority of mitochondrial proteins showing half-life of 16 or greater hours (**Figure 4A**). On the other hand, proteins classified as nuclear or localized to the cell membrane and extracellular space showed significantly faster turnover (ANOVA/Tukey HSD P < 0.05), with a substantial portion showing half-life of 4 or fewer hours (**Figure 4A**). On a pathway level, proteins with greater replacement flux (i.e., shorter half-life) are significantly enriched (Gene set enrichment analysis, FDR < 10%) in extracellular signaling and cell cycle processes (**Figure 4B**), suggesting these processes are associated with active turnover in hiPSC.

**Figure 4.**
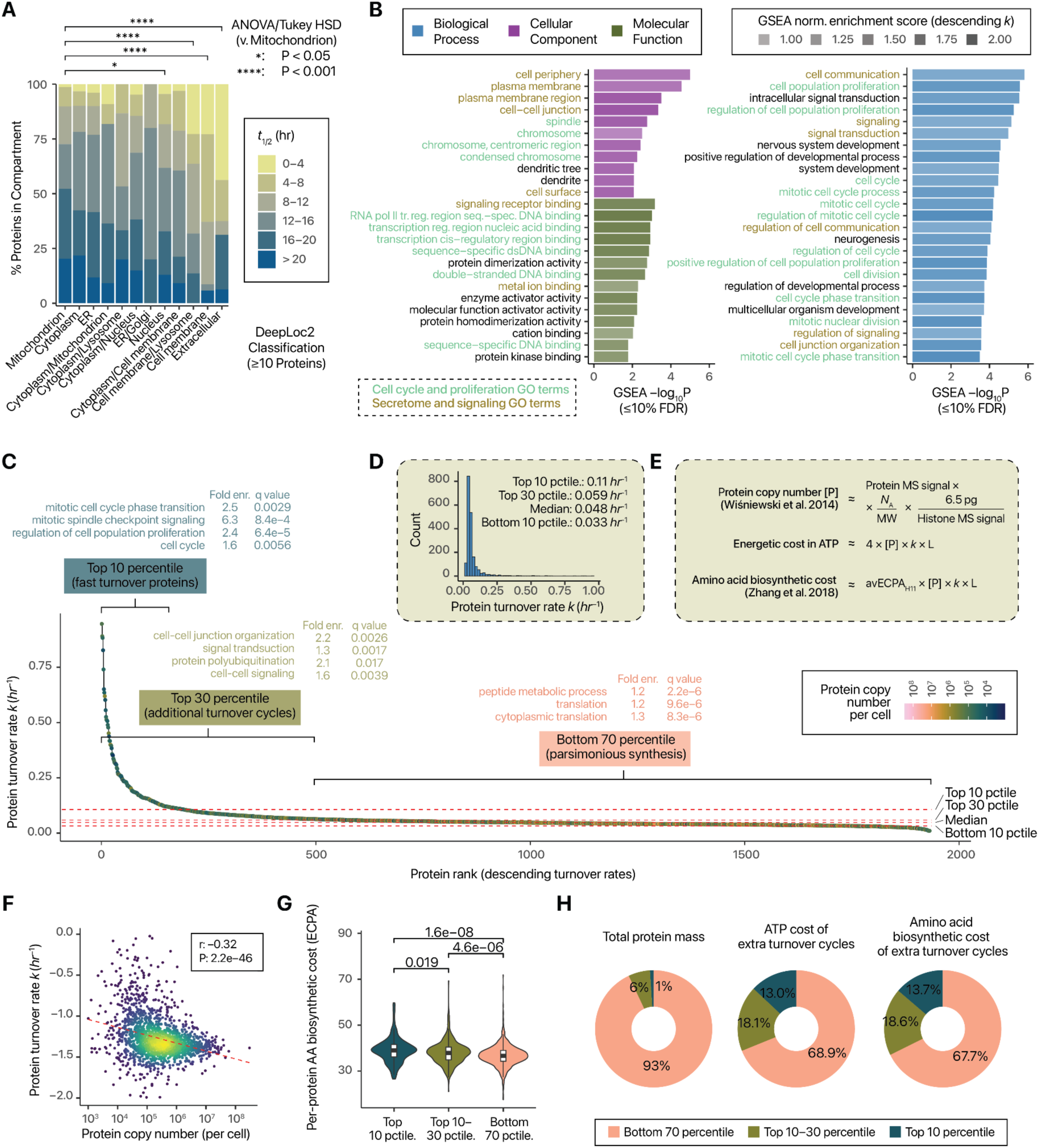
Features of hiPSC proteome-wide turnover dynamics. **A.** Stacked bar charts showing the distribution of proteins half-life values in each of 13 subcellular compartments predicted using DeepLoc2. **B.** GSEA of proteins with descending replacement rates against GO Cellular Component (purple), Molecular Function (olive), and Biological Process (blue) terms. Fast-turnover hiPSC proteins are involved in cell cycle, secretome, and signaling functions. **C.** Ranked protein turnover rate values in hiPSC. Red dashed lines show the turnover rates at 5th, 25th, 75th, and 95th percentiles. The top 5 percentile (i.e. 95–100th percentile) of proteins are enriched in cell cycle proteins. Colors represent estimated absolute copy number per cell (see panel E). **D.** Histogram of turnover rates showing a right skewed distribution. **E.** Calculation of protein copy number using the proteomic ruler method allows calculation of total energetic costs of protein turnover. Biosynthetic cost per amino acid is estimated using the ECPA method. [P]: copy number; *k*: turnover rate; L: protein length. **F.** Scatterplot showing negative correlation between protein copy number and turnover rates. **G.** High turnover proteins are associated with proteins with high protein average amino acid ECPA. **H.** High turnover proteins are associated with disproportional energetics budget based on ATP cost and amino acid biosynthetic cost. The proportional ATP usage and biosynthetic costs of proteins are calculated based on turnover rates beyond what is assumed to be minimally needed for cell doubling, at 90^th^ percentile of global protein turnover rates.

In proliferating cells, a baseline, apparent turnover reflects the contributions of protein synthesis due to turnover and cell growth. A quicker turnover of a protein indicates it is associated with additional protein synthesis-degradation cycles beyond what is required by cell doubling. We therefore further investigated the landscape of proteins showing baseline vs. excess turnover rates hiPSC and the biological or sequence features that may regulate them. We find that the majority of protein turnover rates occupy a relatively narrow band (**Figure 4C**) and that the distribution of protein turnover rates is heavily right-skewed (i.e., right-tailed) (**Figure 4D**). At the bottom 10^th^ percentile, the measured protein turnover rate in hiPSC is 0.033 *hr*^−1^ (**Figure 4D**). This corresponds to the observed effective half-life of 21.2 hours, in line with our observations on hiPSC doubling time and the typical hiPSC doubling time reported in the literature (24–36 hours) ^34^. The turnover rates appear not to follow a log-normal distribution (D’Agostino P: 3.7e–93; skewness 1.54), contrary to observations in other proteomes in vivo ^35,36^. These results reveal that protein synthesis in hiPSC is highly parsimonious, where a considerable majority of proteins are not synthesized and degraded in superfluous turnovers cycles beyond the requirement of cell proliferation. This contrasts with observations in animals, where median protein half-life is typically shorter than DNA half-life (e.g., ∼4 days ^37^ vs. 200+ days in the mouse liver), indicating an average protein pool is turned over many times within a cell in vivo.

Based on the turnover rate distribution, we categorized hiPSC proteins into three groups: proteins in the bottom 70 percentiles with little synthesis beyond needed for cell doubling (half-life > 11.8 hours); proteins in the 10^th^ to 30^th^ percentiles (half-life 6.6 – 11.8 hours), which show appreciable excess turnover cycles; and proteins in the top 10 percentiles (half-life ≤ 6.6 hours), which show rapid excess turnover. Consistent with the proteome-wide gene set enrichment analysis result, fast-turnover proteins are significantly over-represented (Fisher’s exact test q-value < 0.01) in cell cycle and proliferation signaling proteins, whereas mid-turnover proteins are also enriched in signaling and polyubiquitination processes, and slow-turnover proteins are associated with translation and metabolic processes (**Figure 4C; Table S8–S10**). The landscape of protein turnover therefore reflects regulations where proteins associated with different biological functions are synthesized and degraded to different extents.

The production of new protein molecules accounts for a substantial portion of the bioenergetic and biosynthetic budget of proliferating cells ^38,39^ and is constrained by selective pressure ^40^. To explore the energy expenditure of protein synthesis in the hiPSC data, we calculated the absolute copy number value per cell of quantified proteins (Methods; **Figure 4E**). The resulting numbers suggest 4.5% [4.1–4.8%] of histone mass as a total protein mass, which is roughly comparable to previous estimates in cancer cells (∼2–4%). Moreover, this method leads to an estimate of hiPSC cell mass of 826 [736–916] ng (95% c.i.), which is in line with estimates from cell size and proportional protein mass (555–959 pg). Furthermore, the protein copy numbers are consistent across time points and correlate strongly with prior estimates in embryonic mouse fibroblasts ^12^ (Pearson’s r: 0.76 [0.74 – 0.78]; P: 2.2e–313 in log-log scale.) despite differences in methodology (**Figure S3A-B**). These results support that accurate absolute abundance estimates in hiPSC can be derived from D_2_O labeled data.

We observe a robust negative correlation (Pearson’s r: –0.32; P: 2.2e–46) between protein turnover rates and copy number per cell (**Figure 4F**), in other words, positive correlation between abundance and half-life. This is consistent with prior observations ^41^ and may reflect a general constraint that high abundance proteins need to be stable and moreover can only be synthesized at a low rate due to energetics and ribosome capacity limits ^39,42^. We find that the turnover rate is not explainable by simple biochemical features of protein sequences such as isoelectric points, hydropathy, molecular weights, or lengths (**Figure S4A–D**), or large structural categories including intrinsic disorder protein status on DisProt ^43^, helix and strand folds, and the presence of turns (**Figure S5A–D**). The lack of correlation with simple biochemical features is consistent with most proteome wide investigations ^13^ but contrasts with earlier reports from limited numbers of proteins (e.g., ^44^).

The copy number quantification allows us to further calculate the peptide chain elongation ATP usage and amino acid biosynthetic cost ^45^ used in protein synthesis (**Figure 4E**). Surprisingly, fast-turnover proteins in hiPSC have higher average amino acid biosynthetic cost than proteins that are synthesized little beyond required for cell doubling (**Figure 4G**). This is consistent with the notion that proliferating cells avoid de novo synthesis of amino acids ^38^, but contrasts with observations in aging mouse brains where proteins with metabolically expensive amino acids are parsimoniously conserved ^36^. Although the interpretation of sequence features demands caution, as biophysical parameters may be secondarily correlated with protein abundance and thus may not independently determine turnover rates ^41^, the analysis here shows that under culture conditions, hiPSC protein synthesis and usage are not constrained by the biosynthetic costs of amino acids. Indeed, although fast-turnover proteins represent a minor fraction (∼6%) of the total protein mass per hiPSC, they can account for up to one-third of the energy budget needed for protein synthesis beyond the minimal levels required for cell proliferation (**Figure 4H**).

### Hyperdynamic proteins in hiPSC are associated with specific degron motifs

The distribution of turnover rates we observed is consistent with the notion that rapidly proliferating cells in culture can remove proteins via passive dilution to daughter cells or by proteolysis ^6^. Proteins targeted by the latter have been referred to as rapidly-degraded or unstable proteins and are hypothesized to represent dedicated processes controlled by protein stability ^8,9^. Below we refer to them as fast-turnover or hyperdynamic proteins to highlight they are not just inherently unstable but both synthesized and degraded quickly under presumed steady state. Despite previous large-scale resources mapping these proteins in multiple human cell lines ^8,9^; we found a number of previously unknown fast-turnover proteins in hiPSC (**Figure 4C**). We note that while a few non-overlap is due to the removal from consideration of secreted and extracellular proteins in a prior study ^9^ (as these proteins may be diluted into the media rather than degraded; see below), other intracellular proteins that are previously not found to be hyperdynamic in other cell types are clearly present, including KIF22, which functions in spindle formation in mitosis, BUB1B which acts as a mitotic checkpoint protein, and PPP1CC, a nuclear phosphatase that participates in cell division, the nucleolar protein HEATR1, splice factor SRSF5, and up to 125 other proteins (**Table S7**). Therefore the identity of fast-turnover proteins is likely to be highly cell specific and reflecting the biology specific to a cell type.

To investigate the protein degradation mechanisms that drive the additional synthesis-degradation cycles of hyperdynamic proteins in hiPSC, we performed a gene set enrichment analysis against sets of proteins containing sequence motifs targeted for protein degradation (degrons) ^46^. In total, we retrieved 1810 degrons from Degronopedia, and analyzed 105 Degronopedia motifs that appear in at least 10 proteins with quantified turnover rates. We find two degrons that particularly showed a significant association (GSEA adjusted P, FDR < 5%) with higher protein turnover in hiPSC, namely the APC/C KEN box (xKENx), and the SPOP degron ([AVP]x[ST][ST][ST]), whereas a BAG6 type degron (LLLL) also shows nominal enrichment at FDR <10% (**Figure 5A**). The enriched degrons recognize a partially overlapping set of proteins, with a considerable overlap especially between APC/C and SPOP. Multiple cell cycle proteins are included, and are enriched especially among KEN box proteins, including CDC20, CENPF, CDCA2, CDCA5, and others (**Figure 5B**). A notable number of degron-containing hyperdynamic proteins also participate in DNA and RNA binding, DNA repair and stress response (**Figure 5B**). Concordantly, proteins that contain the KEN box or the SPOP degron showed significantly elevated turnover rates in hiPSC than proteins which did not possess either motif, while we did not observe evidence of additive effects (**Figure 5C**). Of interest, the enrichment of the APC/C KEN box contrasts with the lack of statistically significant enrichment of the major APC/C degron D box degron (GSEA adj. P: 0.25) or ABBA degron (GSEA adj. P: 0.65) (**Figure 5A**); likewise, SCF motifs are not significantly enriched, indicating that different cell cycle related ubiquitination processes may be differentially regulated in pluripotent hiPSC.

**Figure 5.**
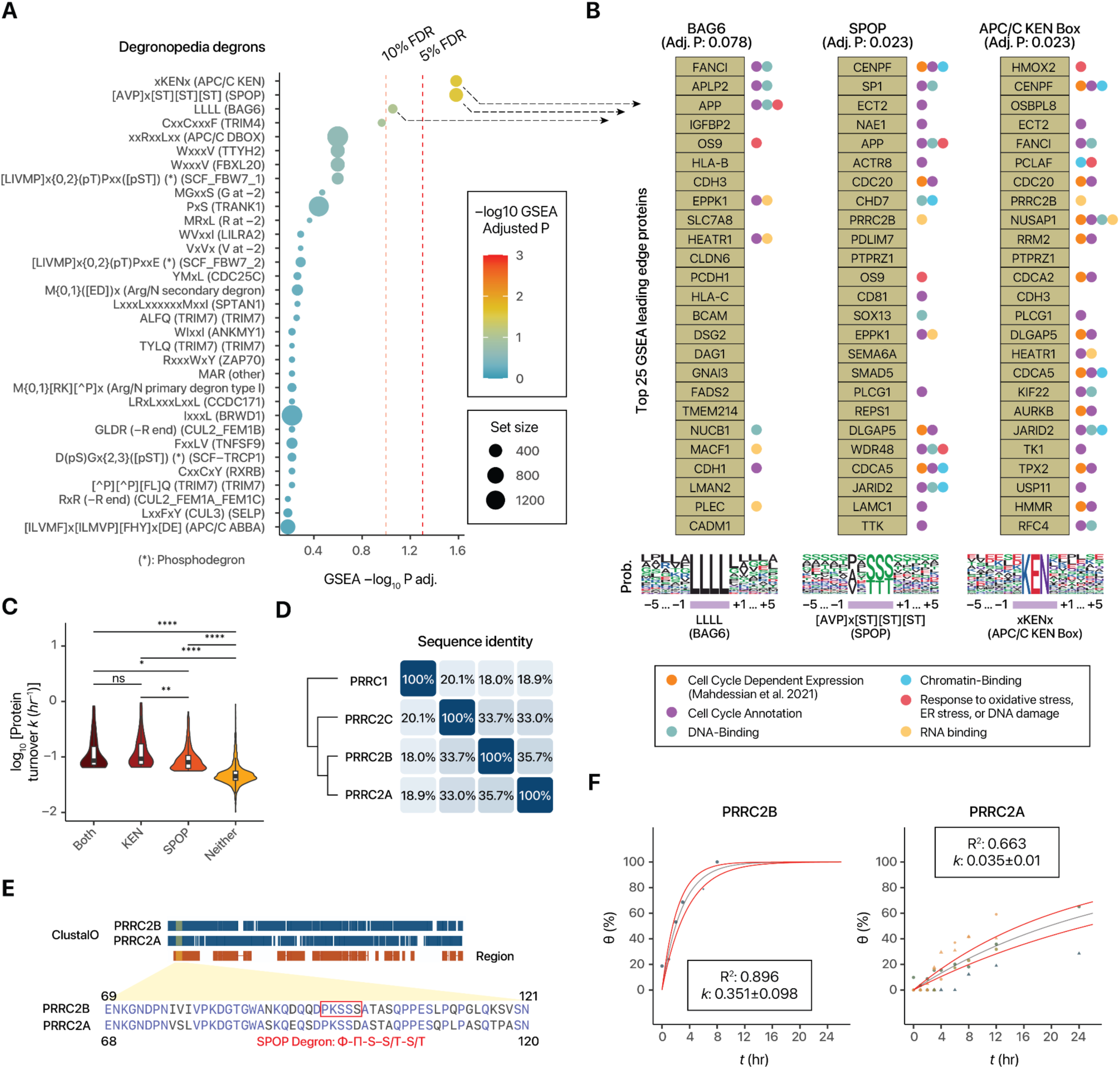
Fast-turnover proteins in hiPSC possess degrons targeted by APC/C and SPOP. **A.** Gene set enrichment analysis (GSEA) of proteins associated with various Degronopedia degrons (top 35 degrons shown), showing the significant enrichment of the APC/C (KEN) and SPOP degrons among fast turnover proteins. **B.** Top 25 GSEA leading edges from the significantly enriched degrons (APC/C KEN, SPOP, and BAG6) are populated by cell cycle dependent, chromatin-binding, DNA-binding, and RNA-binding proteins. **C.** Proteins containing KEN box or SPOP degron have significantly elevated turnover rates in hiPSC. **D.** Sequence identity matrix between PRRC2B and its closest homologs. **E.** SPOP-binding consensus sequence in the PRRC2B N-terminal intrinsically disordered region. **F.** Protein turnover curves fitted from the D_2_O labeling data in hiPSC, showing faster turnover of PRRC2B than PRRC2A.

To investigate how the active protein degradation cycles of dividing hiPSC may be regulated in part by these degradative motifs and activities, we examined PRRC2B/BAT2L/KIAA0515 (Proline-rich coiled-coil protein 2B), a poorly characterized RNA binding protein that exhibits clear excess turnover cycles decoupled from proliferation in hiPSCs (**Figure 5B**). PRRC2B was not previously reported as a fast-turnover protein in four human cell lines from two separate studies ^8,9^, but in our data set is a top GSEA leading edge gene in both the KEN box and SPOP degrons. As PRRC2B is not a known cell cycle protein from annotations or large-scale discovery data ^47^, we focused on the potential effect of the SPOP degron. PRRC2B shares 35.7% sequence identity with its closest homolog PRRC2A/BAT2 (**Figure 5D**), with particularly higher homology in the N-terminal disordered region (residues 1-269 in PRRC2B). Within the N-terminal region, we find a short linear motif (SLiM) ^48^ recognized by SPOP only in PRRC2B (residues 96–100 in PRRC2B; PKSSS) but not PRRC2A (PKSSD) (**Figure 5E**). This sequence also satisfies the more selective definition of the SPOP-binding consensus (SBC) ^49^ motif in the literature (Φ-Π-S–S/T-S/T; Φ: non-polar; Π: polar) and is evolutionarily conserved (**Figure S6**). Elsewhere in the sequence, PRRC2B features three additional SPOP motifs, two of which also satisfy the SBC definition (residues 922–926; 996– 1000), compared to zero in PRRC2A. SPOP (Speckle-type POZ protein) is a substrate-recognizing component of the cullin-RING E3 ubiquitin ligases that localizes to nuclear condensates ^50^. As predicted from the lack of the enriched degrons, we find that PRRC2A does not show strong evidence of elevated turnover in hiPSC (**Figure 5F**). Interestingly, SPOP has been characterized as both oncogenic or tumor suppressive in different contexts, including via the degradation of NANOG ^51–54^. The data here in turn suggests that SPOP-binding sequences might regulate the degradation of multiple hiPSC proteins under baseline conditions, including the participation of PRRC2B in additional turnover cycles but not its homolog PRRC2A.

The APC/C activator CDC20 itself possesses a APC/C KEN box sequence that is known to be targeted by FZR1/Cdh1, as part of a feedback mechanism in the late cell cycle. We note that this KEN box sequence moreover coincides with a predicted PEST motif (epestfind score: 8.85) at residues 97–109 (KENQPENSQTPTK) that is located within a predicted disordered region (MetaPredict ^55^ scores >0.5). Thus, while proteome wide analysis has found no obvious relationship between PEST and turnover ^13,56^, the results here support that PEST motifs may contribute to elevated turnover in specific protein(s) in hiPSC. Prior work has raised the prospect that active protein translation of cell cycle and chromatin regulators is required to maintain a permissive open chromatin, a hallmark of pluripotent mouse embryonic stem cells ^10^. Taken together, the data here indicate that SPOP and APC/C degrons may, at least in part, participate in the physiological synthesis-degradation cycles of proteins that play a role in pluripotency maintenance in hiPSC. More generally, we conclude that D_2_O labeling can be applied to a versatile cell model to measure a wide range of turnover rates, and the resulting data can provide insights into the landscape and regulations of protein turnover cycles in human cells.

### Hyperdynamic iPSC proteins are repressed upon exit of pluripotency

To further explore the roles of hyperdynamic proteins with excess turnover cycles within the context of pluripotency, we re-analyzed data we previously generated on directed differentiation of hiPSC ^57^. Briefly, hiPSC from three donor lines were directed to differentiate using a small molecule based GSK3β inhibition–Wnt inhibition protocol, first toward mesoderm (day 0 to day 2 post differentiation) and then toward cardiac progenitor stage (day 2 to day 6), and protein expression was sampled daily. We then analyzed protein abundance changes during mesoderm and progenitor lineage induction. We find a number of hyperdynamic proteins that exhibit a continued decline in abundance upon differentiation. POU5F1 (OCT3/4) is a hyperdynamic protein and is also one of the master regulators of pluripotency, and its abundance decreased sharply upon directed differentiation as expected (limma FDR adjusted P value < 0.01) (**Figure 6A**). Other hyperdynamic proteins including aurora kinase B (AURKB), which plays a role in chromosome segregation and chromatin remodeling, as well as heme oxygenase 2 (HMOX2), likewise decreased upon differentiation (**Figure 6A**). Moreover, several hyperdynamic proteins overlap with the list of genes required for permissive open chromatin in mouse embryonic stem cells as reported by Bulut-Karslioglu et al. ^10^, including importin subunit alpha-1 (KPNA2/RCH1), elongin-A (ELOA1/TCEB3), and hyaluronan mediated motility receptor (HMMR); which supports that their excess turnover cycle may be associated with their role in maintenance of open chromatin in pluripotent cells, and which becomes lost in pluripotency (**Figure 6B**). On a proteome level, fast turnover proteins have significantly more repressed expression during the 7 days of directed differentiation than proteins without excess turnover cycles (Wilcoxon test P: 5.6e–6) (**Figure 6C**), indicating a number of fast turnover proteins are depleted from hiPSC upon loss of pluripotency.

**Figure 6.**
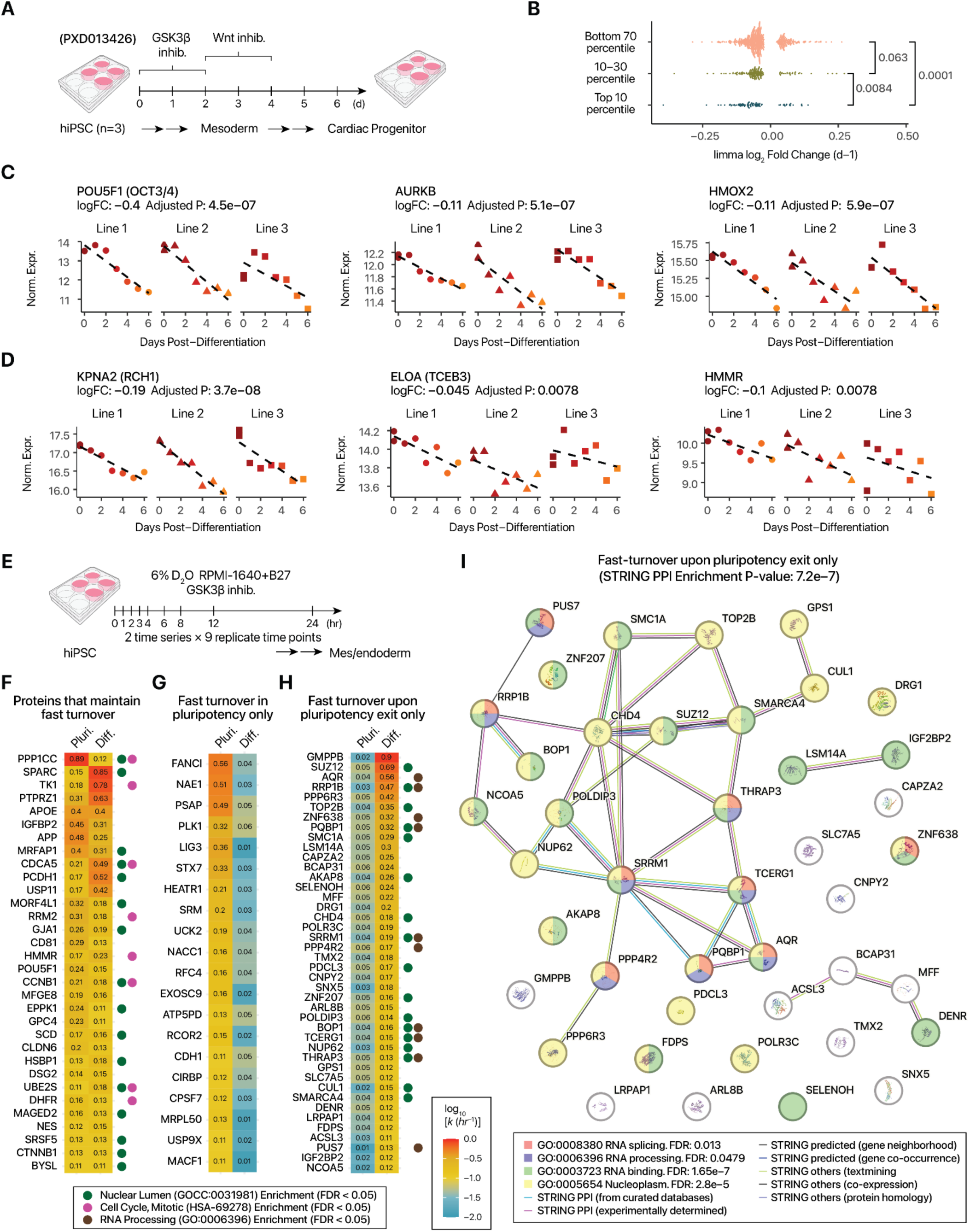
Regulation of fast-turnover hiPSC proteins upon differentiation and loss of pluripotency. **A.** Comparison of fast-turnover hiPSC protein abundance upon loss of pluripotency. Data we previously acquiredfrom 3 lines of hiPSC (PXD013426) are analyzed. **B.** Decline in abundance in log_2_ scale over time post-differentiation (days) upon directed differentiation. Example proteins with significant trends over differentiation (limma adjusted P values < 0.05) are included. Fast-turnover proteins (top 10 percentile) are more likely to decline significantly from hiPSC to mesoderm and progenitor stages. P values: Wilcoxon rank-sum test. **C.** Variance stabilization normalized expression level of three example hyperdynamic genes (POU5F1/OCT3/4, AURKB, and HMOX2) whose expression level decreased upon hiPSC directed differentiation to mesoderm and progenitor stage ^57^. **D.** Same as C, but for three proteins known to maintain open chromatin in mouse pluripotent stem cells (Bulut-Karslioglu et al., 2018) ^10^. **E.** To examine protein turnover changes upon early stages of pluripotency exit, hiPSC is labeled with 6% D_2_O simultaneously with GSK3β inhibition in RPMI-1640+B27 medium for mesendoderm lineage induction. Samples are then collected at 0, 1, 2, 3, 4, 6, 8, 12, and 24 hours post-labeling/post-differentiation. **F.** Heat map of protein turnover rates among proteins that maintain fast turnover cycles (i.e., top 10 percentiles in both pluripotent and differentiating cells). Colors denote log_10_ turnover rates in pluripotent or differentiating cells. Colored circles show proteins associated with two significantly enriched terms (FDR < 0.05) in STRING: nuclear lumen and cell cycle, mitotic. **G.** As in E, but for proteins that become slow-turnover upon differentiation (bottom 70 percentiles in differentiating cells). **H.** As in F, but for proteins that become fast-turnover upon differentiation (from bottom 70 percentiles to top 10 percentiles in the differentiating cell experiment. Colored circles show proteins associated with two significantly enriched terms (FDR < 0.05) in STRING: nuclear lumen and RNA processing. Terms that are not significantly enriched are not labeled. **I.** Gain of new fast-turnover proteins during early mesendoderm induction. STRING network showing a module of proteins which become hyperdynamic within hours of hiPSC directed differentiation. These proteins are significantly enriched in RNA binding, RNA processing, and RNA splicing terms, representing a shift in the biological processes regulated by rapid turnover cycles. Colors in nodes highlight significantly enriched Reactome and Gene Ontology terms over the genomic background. Colors of edges denote STRING network relationship types.

To further examine the fast-turnover hiPSC proteins during the early stages of pluripotency exit, we performed time-course labeling simultaneously with a directed differentiation protocol over 24 hours. Under CHIR-99021 to induce prolonged GSK3β inhibition and Wnt activation, hiPSC exits self-renewal and is directed toward mesendoderm lineage.

Samples are then collected at 0, 1, 2, 3, 4, 6, 8, 12, and 24 hours post-differentiation and post-D_2_O labeling and analyzed by mass spectrometry as the pluripotent cells above (**Figure 6D**), quantifying 6679 unique proteins across all samples and resulting in the peptide-level curve-fitting of 3071 proteins. The cells are now in non-steady-state and moreover we expect some heterogeneity within the differentiating cell population. We previously showed that label incorporation rates are nevertheless approximated by exponential rise to plateau under non-steady-state assumptions ^30^, hence we applied the same kinetic model analysis to estimate the synthesis rates of proteins in differentiating hiPSC, but relaxed the peptide-level R^2^ cutoff to 0.7 and focused on 1,892 well-fitted proteins (**Table S11**). After grouping into fast-turnover (top 10^th^ percentile); moderate turnover (10^th^–30^th^ percentile) and slow turnover (bottom 70^th^ percentile) proteins, the data reveals divergent behaviors of hyperdynamic hiPSC proteins (**Figure 6E**). We find 32 hyperdynamic proteins that are shared between self-renewing and differentiating hiPSC, i.e., in the top 10 percentiles of both data sets, including CDCA5, and CCNB1. These proteins form an interconnected module with more protein-protein interactions than expected over the genome background (STRING P: 8.7e–10), and are significantly enriched in cell cycle and nucleus proteins (**Figure S7**). We find 20 proteins that are hyperdynamic in hiPSC but show low-turnover under differentiation suggesting they are no longer synthesized and degraded in excess upon exit of pluripotency; while these proteins are not significantly enriched in interactions or functional annotations, we note a module of interconnected DNA repair and cell cycle proteins comprising FANCI, PLK1, RFC4, and LIG3 (**Figure 6F**, **S8**), suggesting at least some components of cell cycle regulation are no longer under active proteolytic regulation.

Lastly, a third category of 42 proteins have slow turnover (i.e., bottom 70 percentile) in hiPSC but become hyperdynamic during early differentiation toward mesendoderm. This category is expected to encompass proteins that are undergoing fast synthesis-degradation cycles as well as newly induced proteins under elevated synthesis (**Figure 6G**). Among these proteins are SRRM1, THRAP3, AQR, and PQBPand others that are significantly enriched in RNA binding and RNA splicing function (**Figure 6H**), suggesting protein regulation by active translation-degradation shifts rapidly from primarily regulating cell cycle function toward post-transcriptional processes within hours of pluripotency exit (**Figure 6I**). Taken together, the results support that hyperdynamic proteins are cell type and cell state specific and may represent prime targets for strategies that aim to manipulate pluripotency, lineage determination, and malignancy.

### D_2_O labeling enables secretome flux measurement in hiPSC-derived cardiomyocytes

Lastly, we explored an application of D_2_O labeling to take advantage of fast-turnover proteins to facilitate secreted protein analysis in cell culture. As noted above and also observed in prior work, short-lived proteins in cell culture include extracellular components, which exhibit fast apparent kinetics not necessarily due to proteolytic degradation but because the existing protein pools may be removed during media change, leading to immediate equilibration with background isotope enrichment. We therefore explored the use of D_2_O labeling to facilitate secretome analysis. The secretome plays an important role in cellular communication and crosstalk, but existing mass spectrometry methods are hindered by challenges of distinguishing bona fide secreted proteins from background proteins such as those from culture media supplements (e.g., FBS), basement matrix (e.g., Matrigel) ^58^, or passive leakage due to cell lysis. These problems can be present even where serum-free, albumin-free media is employed^59^. Although methods have been developed that can label secreted proteins metabolically via clickable sugars ^60^, amino acid analogs ^61,62^, or proximity labeling in secretory pathways ^63^, these approaches may either be incompletely bioorthogonal ^62^ or require genetic manipulation in the target cells, and therefore could limit the cell types and subset of secretomes that can be analyzed. Hence, generalizable methods continue to be needed that can distinguish bona fide secretomes in serum free and serum containing media in different cells.

As pre-existing, unlabeled, extracellular proteins are removed during media change and thus newly synthesized proteins are no longer diluted by the pre-existing pools, a sampling of the conditioned media would reveal quick apparent kinetics, which should readily distinguish them from the bulk of high-abundance intracellular proteins that are not synthesized beyond cell doubling requirements. Prior works have also explored the use of dynamic SILAC incorporation to distinguish bona-fide secreted proteins from backgrounds and assess synthesis rates ^64–66^.

We therefore applied D_2_O labeling to analyze cell culture secretomes, focusing on proteins secreted from terminally-differentiated, non-dividing hiPSC-CM. Briefly, hiPSC-CM are cultured in 6% D_2_O enriched RPMI-1640+B27 for 24 hours, upon which the conditioned media is collected and replaced with new 6% D_2_O media; at 48 hours, the media is again collected, and cells are harvested for whole cell lysate. The experimental and analysis workflow is then applied to calculate the fractional synthesis θ and turnover rate *k* of proteins, as well as the flux, calculated as *k* times protein concentration [P] (**Figure 7A**).

**Figure 7.**
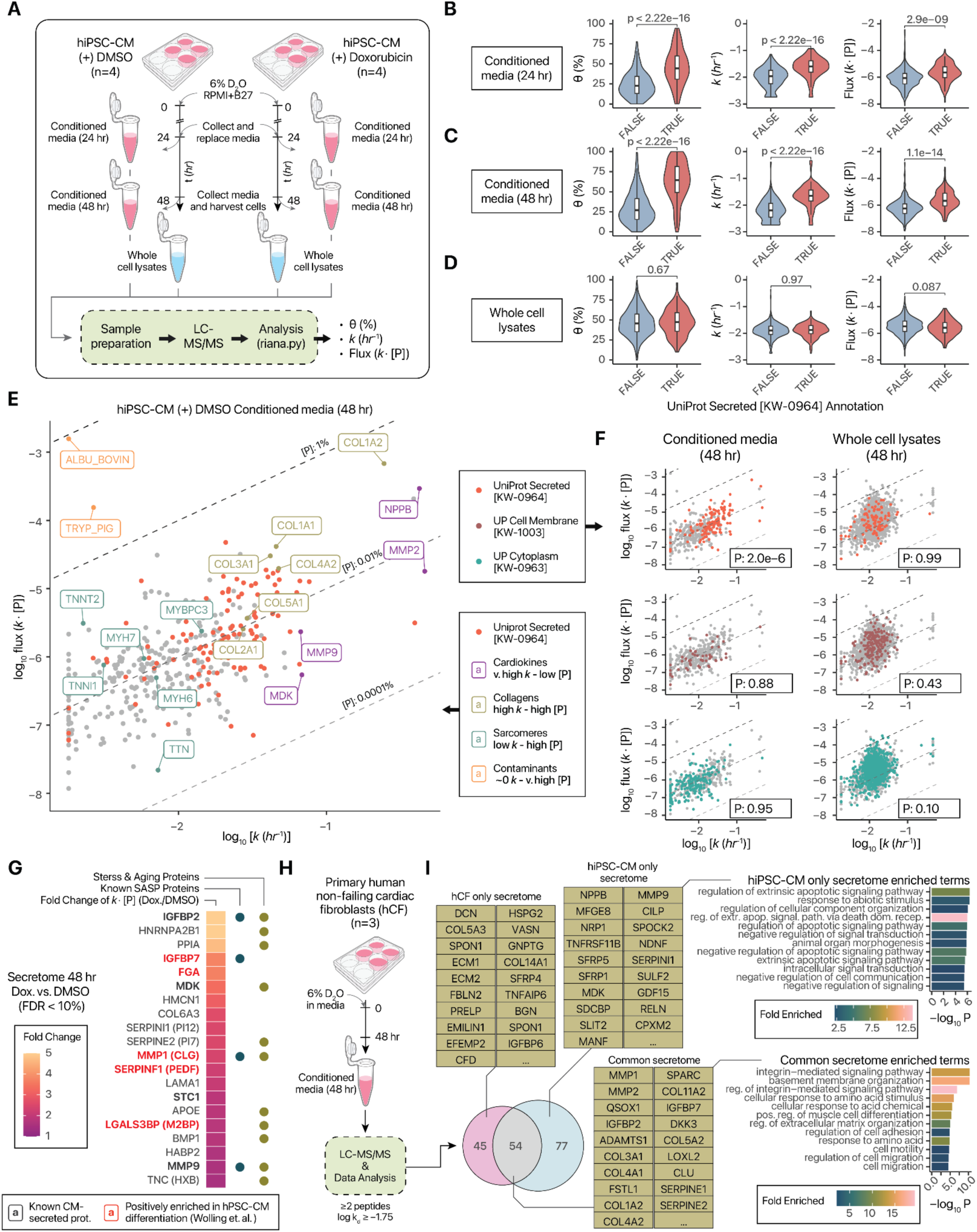
Application of D_2_O labeling to secretome kinetics measurements. **A.** Experimental plans to analyze the secretome of hiPSC-CM treated with doxorubicin or vehicle (DMSO) for 24–48 hours.. **B.** Violin/boxplots (IQR/1.5-IQR) showing the fractional synthesis (left); *k* (center) and flux (right) of proteins in the conditioned media of DMSO-treated hiPSC-CM after 24 hours of label. Proteins are grouped by UniProt annotation for secreted (keyword KW-0964) (blue) vs. non-secreted proteins (red). **C.** Same as B, but for conditioned media collected at 48 hours of labeling. **D.** Same as B, but for whole cell lysates at 48 hours of labeling. **E.** Scatterplot showing the relationship between turnover rates and flux of proteins identified in the conditioned media of DMSO-treated hiPSC-CM at 48 hours of labeling. UniProt Secreted proteins are in red. Colored labels refer to groups of highlighted proteins. **F.** Same as E, but secreted (top), cell membrane (middle), and cytoplasm (bottom) proteins are highlighted for conditioned media (left) and whole cell lysates (right). **G.** Heatmap showing proteins with increases in flux following doxorubicin, colored by fold-change (doxorubicin/DMSO) and annotated with whether they are known CM-secreted proteins, SASP components, or stress and aging related proteins. **H.** Application to analyze the secretome of primary hCF. **I.** Euler diagram showing the overlap in secretome proteins between hCF and hiPSC-CM. Examples of hiPSC-CM-only, hCF-only, and common secreted proteins are shown in tables. Bar charts show significantly enriched GO Biological Processes in hiPSC-CM-only proteins (top) and common secretomes (bottom).

In conditioned media collected after 24 hours, we found that annotated secreted proteins (UniProt KW-0964) showed higher θ, *k*, and flux than non-secreted proteins (**Figure 7B**), a trend that continued in 48-hour conditioned media (**Figure 7C**). Surprisingly, this difference is not apparent in whole cell lysates, which may reflect cell type or experimental design differences from hiPSC (**Figure 7D**). Focusing on the 48-hour conditioned media, we observed that turnover kinetics allowed ready distinction of extrinsic proteins bovine serum albumin (in the culture media) and porcine trypsin (used in protein digestion), both of which have very high [P] but ∼0 *k*, as no deuterium incorporation occurred (**Figure 7E**). Conversely, classically CM-secreted proteins including NPPB, MDK, and MMP2 can be distinguished from intracellular sarcomeric proteins known to leak passively into the surrounding upon cell lysis, as the former have very high *k* and the latter high [P] but low *k*. Next, hiPSC-CM expresses multiple types of collagens ^67^, and we find that collagens have high *k* and [P] in the conditioned media (**Figure 7E**). The fast kinetics of secreted proteins is also reflected in the uneven (left-tailed) distributions of annotated secreted proteins (bootstrap P: 2e–6), contrasted with annotated cell membrane and cytoplasmic/intracellular proteins in conditioned media (**Figure 7F**).

This result corroborates that D_2_O labeling adds discriminant power to distinguish cell secreted proteins from proteins in the media supplement, which may be exploited by heuristics or classification algorithms to identify high-flux secretome components. Using a simple heuristic cutoff of log_10_ *k* ≥ –1.75, which covered ≥ 60% of annotated secreted proteins in the experiments while excluding the highlighted sarcomeric proteins or non-cellular contaminant proteins (4.75-fold OR secreted vs. cytoplasm annotations), we investigated the change of secretome flux in hiPSC-CM at baseline vs. treatment with doxorubicin, a chemotherapy agent that causes cardiac toxicity and senescence-like features. The secretome of doxorubicin-treated iPSC-CM likewise displays telltale kinetics differences between intracellular and secreted proteins (**Figure S9**). Moreover, we observe that doxorubicin treatment induced hiPSC-CM secretion of multiple proteins, including known CM secretomes ^59^, several classical senescence associated secretory phenotype (SASP) components such as IGFBP7 and MMP1 and stress responsive proteins including MDK (FDR <10%) (**Figure 7G**). Thus D_2_O labeling may facilitate the broad identification of perturbation-induced secretome flux in cell models.

We applied D_2_O labeling to primary human cardiac fibroblasts (hCF), obtained from non-failing donor hearts not suitable for transplant, and labeled with D_2_O for 48 hours (**Figure 7H**). Because primary cells are replicatively limited, they can only be cultured for a finite number of passages prior to senescence. We therefore did not pursue a fully labeled calibration standard, and instead opted to apply the deuterium accessible site values from hiPSC-CM toward the analysis. In total, we identified 99 high-flux hCF-secreted proteins and 131 high-flux iPSC-CM proteins passing our criteria, 54 of which were common between two cell types (**Table S12**).

Different classes of proteins are found between CM secreted proteins and commonly secreted proteins. Proteins in cell death regulating and signaling related pathways are statistically overrepresented in hiPSC-CM only secretomes, whereas common and hCF only proteins are significantly enriched in extracellular matrix (ECM) organization and integrin signaling related proteins (**Figure 7I**). Taken together, these results demonstrate that D_2_O labeling can also avail the characterization of cell-specific secretomes within an organ, such as to investigate cellular crosstalk and paracrine signaling.

### Limitations of the Study

There are also potential limitations of D_2_O labeling, compared to other techniques. (1) We assume rapid and constant equilibration rates of D_2_O with amino acids. This is well established in animals, but in cell culture might be influenced by cell feeding, where media change can flood the cells with unlabeled amino acids ^24^. (2) Because of the inherent overlap of isotopomer profiles between unlabeled and labeled peptides, the profile of D_2_O labeling represents only the composite turnover arising from both synthesis from degradation kinetics. (3) The derivation of the fraction of new proteins in D_2_O labeling requires accurate measurement of the relative abundance of mass isotopomers within a peptide isotope envelope, which is sensitive to the spectral accuracy of instrumentation. Previous work in vivo has found generally comparable precision in turnover rates measured from D_2_O and SILAC ^37^, but further work is needed to examine the measurement accuracy and precision systematically in different cell models. (4) There are minor differences in labeling sites across cell types that may be due to cellular biochemistry and media compositions. In slow-growing or non-immortalized cells, it may be infeasible to perform fully-labeled calibration standards, hence sites from other cells may be used that only approximate true labeling extent. Future studies may investigate the mechanisms behind labeling differences. These limitations prompt valuable directions for future development.

## Discussion

Protein turnover is a critical component of gene expression regulation and cellular homeostasis, yet methods for measuring turnover rates in vitro continue to be needed that are scalable and broadly applicable to different cells. To measure protein turnover rates on a large scale, the two common approaches are to apply cycloheximide to inhibit protein synthesis then follow the depletion of existing protein, or alternatively use dynamic stable isotope labeling of amino acids in cell culture (dynamic SILAC or pulsed SILAC) to introduce essential amino acids labeled with multiple heavy stable isotope atom centers (e.g., ^13^C and ^15^N in arginine or lysine) into the cell culture medium for a period of time prior to label equilibration ^12,13,68,69^. The incorporation of labels into newly synthesized proteins over time is then quantified with mass spectrometry-based proteomics to provide information on turnover kinetics. Despite their utility, these techniques have limitations that call for the continued development of complementary methods. Cycloheximide treatment can lead to unwanted effects on cellular physiology. Dynamic SILAC is constrained by the high cost of synthetically labeled amino acids, especially where prolonged or large-scale labeling is needed; the inability to label peptide sequences lacking the targeted amino acid(s); and the need to modify culture media to use dialyzed serum and/or amino acid-depleted basal media, all of which can limit its scalability and applicability in certain experimental designs. To illustrate, protein half-life in some primary cells has been shown to reach 50 days or more, approaching in vivo lifetimes of proteins in animal models ^69^. In this scenario, upon 48 hours of SILAC labeling, a slow-turnover protein would have accumulated less than 1% of label, which could complicate accurate quantification and contribute to substantial variances in the calculated half-life, thus demanding prolonged labeling up to a week that could add considerable reagent costs. Other experimental designs, such as those requiring the fractionation of subcellular components ^32^, may require upward of 10^7^–10^8^ cells per experimental condition and could strain the scalability and accessibility of SILAC labeling.

Recent-year methodological advances on analytical frameworks ^28,70^, as well as bioinformatics tools that enable automated data processing on a proteome scale ^35,37,71,72^, have further improved the feasibility of D_2_O labeling studies in animal studies and now enable proteome-wide surveys of protein turnover kinetics ^30,56,73,74^ that are compatible to or even outperform SILAC labeling in animals ^37^. Here, we describe an updated method to perform proteome-wide turnover kinetics studies in cell culture using low-dose D_2_O medium dilution, which should be widely compatible with different culture medium formulations. Previous use of D_2_O to measure protein turnover kinetics in cell culture has been severely limited, and largely relied on picking out individual alanine/glycine rich peptides, and assuming labeling was limited to alanine and glycine and to an identical extent as in adult animals (i.e., 4 deuterium atoms in alanine and 2 in glycine) ^25^. Addressing this critical gap, we determined the deuterium-accessible atoms on all 20 proteinogenic amino acids across multiple cell lines. By resolving the number of labeling sites in proteinogenic amino acids in cell cultures, we demonstrate a workflow to acquire fractional protein synthesis values needed to calculate turnover rates from virtually any peptide in D_2_O-labeled proteomics data. Compared to dynamic SILAC, D_2_O offers the advantages of low cost, convenience, and compatibility with different cell types and cell culture media, and may help facilitate experimental designs that require extended labeling period or scaling up to many cells. As we show that D_2_O labels multiple non-essential amino acids including Ala, Glu, Gly, and Asp, it has the potential to be compatible with wider selection of peptide sequences over common isotope-labeled amino acid tracers, and thus include additional peptides in analyses especially where alternative proteases such as chymotrypsin are used. Moreover, the continued development and applications of next-generation pluripotent stem cell and organoid models also rely heavily on commercial media and differentiation kits for batch consistency and reproducibility, which can be easily compatible with D_2_O labeling as opposed to the need to formulate multiple custom SILAC media.

Taking advantage of the flexibility of D_2_O labeling, we quantified the absolute turnover kinetics of proteins in hiPSC. hiPSC are proliferative cells that maintain open chromatin and a high gene expression entropy. Previous work in mouse embryonic stem cells has shown that active protein translation is needed to maintain open chromatin, whereas cycloheximide inhibition of translation leads to the depletion of cell cycle regulation and chromatin remodeling proteins ^10^. However, it remained unclear to what extent pluripotent cells maintain excess protein turnover cycles beyond proliferative requirements in untreated wild-type cells. Our data confirm a highly parsimonious protein turnover landscape in hiPSC where the majority of hiPSC proteins are not synthesized beyond cell doubling demand and are thus primarily removed via passive dilution to daughter cells. On an organellar level, mitochondrial proteins have particularly low turnover, consistent with hiPSC having high glycolytic rate while limiting the use of mitochondrial oxidative metabolism to maintain pluripotency ^75^.

On the other hand, “hyperdynamic” proteins that turn over independent of cell doubling are enriched in cell cycle regulation, DNA repair, and stress response, highlighting a number of proteins within these processes that may be important for the maintenance of pluripotency and maintaining genome and proteome integrity. Unlike cycloheximide-based approaches, the time-course experiments here allow the timing and kinetics of degradation to be quantified under untreated baseline conditions. We find that some fast-turnover proteins have half-life as short as 1–3 hours, which would indicate they are only synthesized on demand within a particular phase of the cell cycle, or alternatively are synthesized and degraded over many cycles or in large excess. These proteins include many that are not previously known to be short-lived, are enriched in specific degron motifs, and are depleted upon loss of hiPSC pluripotency. These proteins therefore present potential targets for manipulating pluripotency related processes via protein degradation. Moreover, our observations in differentiating cells are entirely consistent with earlier work in mouse embryonic stem cell undirected differentiation, which showed that many nuclear proteins are regulated by translational or translational mechanisms upon cell fate transition (i.e., their protein level changes were not well mirrored by RNA changes) ^76^. The shift in the turnover kinetics of post-transcriptional regulators suggests a mechanism of rapidly rewiring gene expression networks in differentiating cells that merit further exploration.

Extending our investigation of fast-turnover proteins, we adopted D_2_O isotope labeling to analyze cellular secretomes and monitor the kinetics of protein secretion. The inclusion of label kinetics facilitates the determination of bona fide secretome candidates with empirical data, as well as allows total secretion flux to be compared across conditions. Such kinetic information may be important for determining the biological nature and contexts of protein secretion events. A high secretory flux may indicate a specific crosstalk reaction in response to stimuli, which may help detect secreted biomarkers, such as those associated with chemotherapy-associated cardiac toxicities. On the other hand, delayed kinetics may reflect stalling across secretory pathways, and is being targeted in experimental cancer therapy to reduce release of tumor-associated proteins.

In summary, we describe a simple and accessible method for isotope incorporation kinetics experiments in cultured cells, by simply adding low-volume D_2_O labeling to standard tissue culture media. D_2_O labels primarily non-essential amino acids not supplemented in media to enable mass spectrometry analysis of isotope incorporation rates. This allows accurate interpretation of D_2_O labeled mass spectra to extract protein kinetics information, and enables proteome-scale turnover measurements from single or multi time-point continuous labeling designs. We foresee this method will broadly enable research efforts to study protein homeostasis regulations in cells and find applicability in multiple areas.

## Acknowledgments

We thank Dr. R. J. Beynon at the University of Liverpool for helpful discussions and insights. We thank Drs. M. R. Bristow, J. A. Schwisow and E. Jonas for assistance with procuring human heart samples; and Drs. P. Buttrick, A. Ambardekar, B. Kopecky and the University of Colorado’s Division of Cardiology for ongoing maintenance of the human cardiac tissue biobank. This work was supported in part by National Heart, Lung and Blood Institute (NHLBI) grant R01HL171711 and an American Heart Association Collaborative Sciences Award (24CSA1255857) to T.A.M.; National Institute of General Medical Sciences (NIGMS) award R35GM146815 and NHLBI grant R00HL144829 to E.L; NHLBI grant R01HL141278, NIGMS grant R01GM144456, and the University of Colorado SOM Advanced Proteomics Infrastructure (API) funds to M.P.L. J.G.T. was supported by an NHLBI Pathway to Independence Award K99/R00HL166708. The content is solely the responsibility of the authors and does not necessarily represent the official views of the National Institutes of Health.

## Author Contributions

Conceptualization, M.P.L and E.L.; Methodology, M.P.L. and E.L.; Investigation, L.A., D.C.N., J.C., A.B., B.P., J.P., P.S., and J.G.T.; Writing – Original Draft, M.P.L., and E.L.; Writing – Review & Editing, M.P.L. and E.L.; Funding Acquisition, T.A.M., M.P.L. and E.L.; Resources, T.A.M., M.P.L., and E.L.; Supervision, M.P.L. and E.L.

## Methods

### Human AC16 cell labeling calibration standards

The AC16 calibration standards data were initially generated for our parallel work to evaluate the effect of isotopomer selection on fractional synthesis calculation ^28^, and here utilized for an independent study goal. Briefly, AC16 cells (Millipore) were cultured in monolayer on tissue culture plates with DMEM/F12 medium (Gibco) supplemented with 10% FBS. The basal medium was diluted with either 6% D_2_O (heavy labeled population) or 6% H_2_O (control population) at 37°C, 5% CO_2_. The cells were maintained in this medium for 3 passages, each passage with a split ratio of 1:8. This growth was estimated to constitute approximately 9 doublings of the cell populations. The cells were harvested by trypsinization, pelleted, washed once with phosphate buffered saline, and pelleted again before snap freezing in liquid nitrogen and storing at –80°C. At the time of processing each pellet was resuspended in 1 mL of RIPA buffer (Thermo Scientific) supplemented with Halt Protease and Phosphatase Inhibitor Cocktail (Thermo Scientific). Proteins were extracted with sonication in a Bioruptor Pico (Diagenode) with settings 10× 30 sec on 30 sec off at 4°C. Insoluble debris was pelleted and removed from all samples by centrifugation at 14,000 ×g, 5 minutes.

Protein concentration of all samples was measured with Rapid Gold BCA (Pierce). Cell lysates from the D_2_O and H_2_O media populations were then combined in a labelling series expressed as the proportion of protein that was labeled with heavy water: 0, 0.125, 0.25, 0.375, 0.5, 0.625, 0.75, 0.875 and 1. The samples were trypsin digested using a modified version of the filter-aided sample preparation approach as previously described. A total of 50 µg protein per sample in 250 µL 8 M urea was loaded onto Pierce Protein Concentrators PES, 10K MWCO (Thermo Scientific) pre-washed with 100 mM ammonium bicarbonate (ABC). The samples were again washed with 8 M urea to denature proteins and remove SDS. The samples were washed with 300 µL 100 mM ABC twice. The samples were then reduced and alkylated with final concentrations 5 mM dithiothreitol (DTT) and 18 mM iodoacetamide (IAA) for 30 minutes at 37°C in the dark. DTT and IAA were removed with centrifugation and the samples were washed 3× with 100 mM ABC. Samples were digested atop the filters overnight at 37°C with mass spectrometry grade trypsin (Promega) at a ratio of 1:50 enzyme:protein. The following day samples were cleaned with Pierce C18 spin columns (Thermo Scientific) according to the manufacturer’s protocol. Eluted peptides were dried under vacuum and redissolved resuspended in 0.1% (v/v) formic acid.

The samples were analyzed on a Thermo Q-Exactive HF quadrupole-Orbitrap mass spectrometer coupled to a nanoflow Easy-nLC UPLC with the Thermo EasySpray electrospray ionization source. Peptides were separated with a PepMap RSLC C18 column 75 μm × 15 cm, 3 μm particle size (Thermo Scientific) with a 90 minute gradient from 0 to 100% pH 2 solvent B (0.1% formic acid in 80% v/v LC-MS grade acetonitrile). The mass spectrometer was operated in data-dependent acquisition (DDA) mode with scans between m/z 200 and 1650 acquired at a mass resolution of 60,000. The maximum injection time was 20 ms, and the automatic gain control was set to 3e6. MS2 scans of the 15 most intense precursor ions with charge states of 2+ to 5+ were acquired with an isolation window of 2 m/z units, maximum injection time 110 ms, and automatic gain control of 2e5. Fragmentation of the peptides was by stepped normalized collision-induced dissociation energy (NCE) of 25 to 27. Dynamic exclusion of m/z values was used with an exclusion time of 30 seconds. The DDA mass spectrometry data were searched against UniProt Swiss-Prot database ^77^ retrieved using Philosopher v.4.8.1 ^78^ on 2023-06-27 with added contaminants using Comet v.2022.01 ^42^ including the following parameters: decoy_search = 1; peptide_mass_tolerance: 10.00 ppm; num_enzyme_termini = 1; isotope error: 0/1/2/3; fragment_bin_tol = 0.02; fragment_bin_offset = 0.0. Search results were post-processed using Percolator (crux-4.1 distribution) ^80^ with the following options: –-decoy-prefix DECOY_; –-overwrite T; –-maxiter 10. Peptide identifications at FDR adjusted q value of greater than 0.01 were excluded. Isotopomer intensity of the m_0_, m_1_, m_2_, m_3_, m_4_, and m_5_ peaks was extracted using Riana v.0.8.0 integrate ^37^ with the following settings: –q (q-value) 0.01, –r (retention time in minutes) 0.25 –m (mass tolerance in ppm) 15.

### Fractional synthesis calibration cells for hiPSC and hiPSC-CM

New calibration standards for hiPSC and hiPSC-CM were generated in this study. Briefly, AICS-0052-003 hiPSC (mono-allelic C-terminus mEGFP-tagged MYL7 WTC-11; Allen Institute Cell Collection) line was acquired from Coriell Institute and seeded onto Geltrex (Gibco) coated 6 well plates and maintained in mTesR Plus basal media with Supplement (STEMCELL) media.

The basal medium but not supplement was diluted with either 6% D_2_O (heavy labeled population) or 6% H_2_O (control population), and the cells were cultured at 37 °C, 5% CO_2_ with daily media changes. At 80% confluency, cells were passaged at 1:6 using EDTA before resuspension in media supplemented with 10 µM Y-27632 (Selleck), for a total of 4 passages (equivalent to ≥10 doublings). The cells were then harvested by EDTA, processed, and analyzed by mass spectrometry as above.

To produce calibration cells for hiPSC-CM, fully labeled or unlabeled hiPSC were replated into Geltrex coated 12 well plates at a density of 4.5 × 10^5^ cells/well and daily media changes continued until the cells reached 80% confluency, day 0 of cardiac differentiation. The hiPSC were differentiated into hiPSC-CM using a small molecule-based GSK-3β inhibition/Wnt inhibition protocol ^81^. Briefly, on day 0, cell media was replaced with 2 mL/well RPMI supplemented with B-27 minus insulin (Gibco) and 6 μM CHIR99021 (STEMCELL); on day 2, the media was changed to 2 mL/well RPMI+B-27 minus insulin. On day 3, the media was changed to 2 mL/well RPMI+B27 without insulin supplemented with 5 μM IWR-1-Endo (STEMCELL). On day 7, the media was changed to 2 mL RPMI+B27 with insulin. Differentiation was confirmed via visualization of morphology, spontaneous contraction of cells, and imaging of the GFP tagged MYL7/MLC-2a. Media was refreshed every other day with RPMI+B27 with insulin until day 14. Attached hiPSC-CMs were washed twice with PBS before incubating with 0.5 mL per well of TrypLE Express (1x, Thermo Fisher) for 11 minutes. An additional 1.5 mL PBS was added to dilute the TrypLE and the cells were triturated until fully detached. Cells were pelleted by centrifugation at 300 ×g for 3 minutes, washed with PBS before repelleting, and stored at –80°C until protein extraction. The samples were then processed and analyzed by mass spectrometry as above.

### Prediction of deuterium labeling sites in cell culture

Because D_2_O labeling typically uses enrichment of 10% of lower excess deuterium, the resulting isotope pattern of labeled proteins is complex and overlaps with the unlabeled protein.

Moreover, whereas in SILAC labeled amino acids are pre-synthesized with fixed mass shifts, upon D_2_O labeling, each amino acid residue has a characteristic number of deuterium-accessible labeling sites based on their biochemistry, hence the extent of mass shift a peptide exhibits upon labeling is sequence-dependent. Analysis therefore requires knowing the number of deuterium-accessible labeling sites in each amino acid, *S*_aa_. This information may be learned from asymptotically labeled peptides or nested fitting over many peptides, but this is not always practical to achieve (e.g., in short-term labeling or labeling of only few time points) and may accrue additional greater fitting error, therefore calling for a general method to calculate the fractional synthesis of any peptide sequence irrespective of label duration.

To recover the fraction of newly synthesized protein *θ* (also referred to as *f* in the literature. We avoid the use of *f* due to potential confusion with other fractional or functional notations), isotope incorporation is analyzed as the change in the ratio between the monoisotopic peak over the complete isotope profile (*A*_0_) across labeling time points (or in the case of the calibration experiment, across known experimental proportions of ground-truth *θ*). As fractional synthesis 0 ≤ *θ* ≤ 1 increases, *A*_0_(*θ*) decreases linearly from the initial position, toward the asymptote as determined by the excess deuterium enrichment (*p*) and number of accessible labeling sites of the peptides, *S*_pep_ (sometimes denoted as *n* or *N*_EH_ in the literature; *S*_pep_ is used here to avoid confusion with number of replicates or the number of amino acids per peptide.)

The initial A_0_ (i.e., unlabeled, *A*_0_(*θ*=0)) of a peptide can be calculated from natural isotopic distribution. The fully labeled *A*_0_(*θ*=1) of the peptide is calculated from the naturally occurring A_0_, the total number of deuterium exchange sites on the peptide *S*_pep_, and the deuterium relative isotope abundance *p* where *p* = 0.06 in the 6% D_2_O experiments:

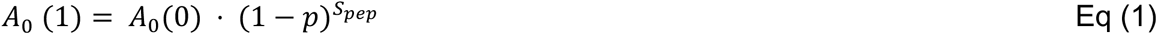

This could be refined by considering naturally occurring deuterium ^82^, but the background deuterium level (0.0001157098) is negligible compared to *p* and is ignored here.

The number of deuterium accessible labeling sites in each amino acid, *S*_aa,_ in human cell culture were predicted by two machine learning strategies. In the first method, a linear regression model in is used to find *S*_aa_:

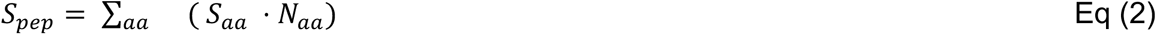

Peptide series from the calibration standards that were identified at Percolator FDR 1% and quantified in all 9 mixture proportions were included. The peptides were then further filtered to include those with a linear fit (R^2^ > 0.9) of mixture proportion *θ* against *A*_0_. Training is done using the LinearRegression model in scikit-learn ^83^, with the settings fit_intercept=False, positive=True. following 80/20 train-test split with 1337 peptides in the training set and 335 in the test set.

In the second strategy (direct prediction of isotopomer profiles), the empirical isotopic ratios as measured in the 25.0%, 50.0%, 75.0%, and 100.0% proportion of fully labeled cell lysates were used as targets of training. The full isotopomer profile of an unlabeled peptide is calculated using the isotope fine structure calculation algorithm IsoSpec2 ^29^ to resolve exact isotopologues of a peptide given its chemical composition. The isotope envelope of a labeled peptide is then approximated by performing the calculation with a modified elementary composition table to include a new element H* (exchangeable hydrogen) with 1 – *p* probability of having mass 1.007825, and *p* probability of having mass 2.014102. A train-test split of test size 0.2 was performed, resulting in 7,200 samples (peptide–proportion combination) in the train sets and 1,800 samples in the test sets. We then used the differential evolution algorithm in scipy.optimize ^84^ to perform global optimization varying the labeling site of each amino acid residue as input to IsoSpec2 and then minimizing the median absolute scaled errors (m.a.s.e., calculated as median absolute error divided by median absolute deviation) between the IsoSpec2 predicted isotopic cluster and the actual empirical ratios of m_0_/m_1_, m_0_/m_2_, m_1_/m_2_, and m_1_/m_3_ in each of the four mixture proportion experiments. The differential evolution parameters are tol: 1e-5, disp: True, polish: False, maxiter: 2500, seed: 42. Lower and upper bounds of labeling sites were set between 0.08 and the number of hydrogen atoms per amino acid residue, respectively.

### Acid hydrolysis of AC16 lysate and direct infusion mass spectrometry

To experimentally validate the predicted labeling sites in cell culture, non-labeled (i.e., 0% D_2_O) and fully labeled (i.e., nine doublings under 6% D_2_O) human AC16 cell lysates (∼200 µg) were desalted and cleaned up using Zeba desalting columns (Thermo) with 7 kDa molecular weight cutoffs (MWCO). The samples were then acid hydrolyzed with 1 mL of 6 M of HCl at 110 °C for 48 hours in a vacuum hydrolysis tube filled with N_2_ to prevent oxidation. The hydrolysate was cooled under a steady stream of N_2_ in a water bath at 90 °C and then dried using a SpeedVac evaporator (Thermo). The dried samples were reconstituted with 300 µL of methanol/water (50:50) with 1% *v*/*v* formic acid, and directly infused with a syringe into a Q-Exactive HF Orbitrap mass spectrometer through a Nanospray Flex ion source. The following mass spectrometry settings were used to acquire MS1 signal for the hydrolyzed amino acids: ESI voltage of +3 kV, injection flow rate of 3 µL/min, mass range of m/z 50–250, scan time of 1 min, FT resolution of 30,000, 60,000, and 120,000, capillary temperature of 275 °C, AGC Target of 1e6, max ion time of 50 ms, micro scan of 1 count.

### AC16 and hiPSC culture for dynamic D^2^O labeling experiments

For the baseline and stressed AC16 cells, AC16 cells were grown in monolayer as above. While in log phase growth, cells were introduced to 6% D_2_O with or without 1 µM thapsigargin for 16 to 24 hours. For hiPSC, cells were labeled in 6% D_2_O in mTeSR Plus and then collected at 9 time points at 0, 1, 2, 3, 4, 6, 8, 12, and 24 hours after the start of labeling. The cells were then harvested, and processed for DDA mass spectrometry analysis as described above.

For hiPSC time-course labeling, AICS-0052-003 hiPSC were passaged as above. Once the cells reach 50% confluency, mTeSR Plus media diluted with 6% D_2_O (heavy labeled population) or 6% H_2_O (control population) was introduced to the cells for the duration of each time interval at 0, 1, 2, 3, 4, 6, 8, 12 and 24 hours as described in the text. At the end of each time interval, the labeling medium was aspirated, the cells were washed with PBS, and then collected with dissociation media (0.5 mL EDTA in PBS), and snap-frozen immediately. Proteins were then extracted using RIPA buffer with BioRuptor sonication as described above, then digested and analyzed with DDA mass spectrometry.

### Directed differentiation of hiPSC for dynamic D^2^O labeling experiments

For hiPSC differentiation toward mesoderm/mesendoderm specification, ACIS-52 hiPSC were plated onto a 6-well plate coated with geltrex in mTesR Plus basal media with Supplement (STEMCELL) media and 10 µM Y-27632 (Selleck). Media changes were performed every day with mTesR Plus basal media with Supplement (STEMCELL) media. At 80% confluency, cells were split with TrypLE Select Enzyme (10X, Thermo Fisher) onto geltrex coated 12-well plates at 0.2 × 10^6^ cells/well in mTesR Plus basal media with Supplement (STEMCELL) media and 10 µM Y-27632 (Selleck). Media changes were performed every day with mTesR Plus basal media with Supplement (STEMCELL) media. At 80% confluency, the medium was removed and replaced with 2 mL/well RPMI1640 supplemented with B-27 minus insulin (Gibco), 6 μM CHIR99021 (STEMCELL), and 6% v/v D_2_O (Cambridge Isotope), except for time point 0. At each timepoint of 0, 1, 2, 3, 4, 6, 8, 12 and 24 hours, media was aspirated from cells and cells were collected with 100 µL TrypLE Select Enzyme (10X, Thermo Fisher) for 2 minutes, then quenched with 500 µL RPMI1640. For each replicate in each condition, 2 wells were combined, pelleted at 200 ×g for 5 minutes, rinsed 1× with 1 mL PBS, pelleted once more at 200 ×g for 5 minutes and immediately flash frozen. Protein extraction, digestion and peptide desalting were performed using the same protocol described above.

The digested peptides were resuspended in 0.1% formic acid (v/v) and separated using a Vanquish Neo UHPLC system (Thermo Fisher Scientific) on an Easy-Spray™ analytical column (ES900 C18, 75µm x 150 mm, 3 µm) with a 60-minute gradient. Solvent A was 0.1% formic acid in LCMS-grade water and solvent B was 0.1% Formic acid in 80% LCMS-grade acetonitrile. The LC-system was coupled to an Orbitrap Exploris 480 (Thermo Fisher Scientific) operating in positive mode with the spray voltage at 2 kV and ion transfer tube temperature of 275 °C. Full-mass spectra were acquired from 350–1650 m/z at 60,000 resolution with an AGC target at 300% and maximum ion injection time set to auto. The mass spectrometer was operated in DDA mode, selecting precursors within a charge state of 2–6 and minimum intensity of 3 x 10^2^. The top 20 precursors were subsequently selected with a 2 m/z isolation window for HCD fragmentation with NCE set at 30%. The resolution was set to 15,000 with an AGC target at 200%, maximum ion injection time of 26 ms and dynamic exclusion of 45 seconds. The data were then analyzed as above.

### Protein turnover kinetics analysis of mass spectrometry data

Isotopomer intensity was extracted using Riana v.0.8.0 to extract the intensity over time of the m_0_, m_1_, m_2_, m_3_, m_4_, and m_5_ peaks as described above. For each peptide, S_pep_ is calculated from the predicted S_aa_ values (**Table S1**) using Eq (2). The plateau enrichment of a peptide can then be predicted using a modified version of Eq (1), that is, for *A*_0_ (*t* = ∞). The fractional synthesis of a peptide at time *t* can then be calculated as:

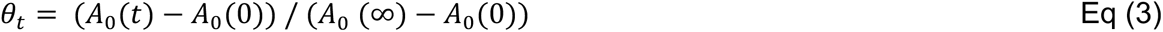

The time series of fractional synthesis at one or more experimental time points was then fitted to a simple exponential kinetic model (Eq (4)) to obtain the best-fit turnover rate constant (*k*) to explain the time series, using the optim function in R.

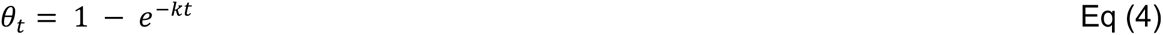

For AC16 single time-point labeling, no goodness-of-fit may be acquired. Therefore the best-fit turnover rates of the top 6 peptides ranked by intensity were used to calculate the median as the protein-level turnover rate, and proteins quantified with at least 3 peptides and with a relative median absolute deviation (calculated as median absolute deviation divided by median of the turnover rates) of 33% or less were accepted. For time-course labeling, goodness-of-fit is determined by the residual sum of squares (RSS) and total sum of squares (TSS) of the kinetic model curve-fitting:

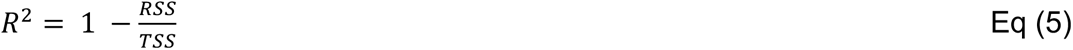

A peptide-level R^2^ ≥ 0.8 is accepted as described in the text. All fractional synthesis data for all constituent peptides that passed this filter are then collected for a single protein-level fitting, where the sums of square weighted to a normalized log intensity of the peptide are minimized by the optim function in R. A protein-level weighted R^2^ ≥ 0.8 is accepted as described in the text.

### Secretome analysis in D_2_O labeled hiPSC-CM

AICS-0052-003 hiPSC (mono-allelic C-terminus mEGFP-tagged MYL7 WTC-11; Allen Institute Cell Collection) were expanded and differentiated into hiPSC-CM as above. Media was refreshed every other day with RPMI+B27 with insulin until day 41, at which time the media was supplemented with 6% *v*/*v* D_2_O (Cambridge Isotope), and 1 µM doxorubicin or vehicle. 24 hours later, the media was collected and replaced with media supplemented with 6% vol/vol D_2_O. The collected conditioned media was centrifuged for 5 minutes at 300 ×g and 5 minutes at 14,000

×g. Spun media was stored at –80°C. 24 hours later (48 hours of labeling total), the media was collected again and centrifuged as described. At the same time, the intracellular samples were collected and reduced, alkylated, and digested using an on-filter digestion protocol as described above.

To analyze secretome content, 300 µL of spun conditioned media per sample was depleted using the Seer Proteograph XT workflow (Seer, Inc.) utilizing two distinct nanoparticle (NP) mixtures (NPA, NPB). NP protein coronas were reduced, alkylated, and digested with Trypsin/Lys-C to generate tryptic peptides for LC-MS analysis. Digested peptides were desalted, eluted in a high-organic buffer into a deep-well collection plate, and quantified. In hiPSC-CM experiments, cleaned peptides were reconstituted in 0.1% formic acid to a final concentration of 0.0325 µg/µL and 0.116 µg/µL from NPA and NPB, respectively. 3 µL of peptides per sample were separated with online low-pH reversed-phase LC (PepMap C18 column, 3-μm particle, 100-Å pore; 75 µm × 15 cm; Thermo Fisher Scientific) via the EASYnLC 1200 system coupled to the Easy-Spray ion source (Thermo Fisher Scientific) at 300 nL/min with a 90-min gradient: 0–75 min: 0 to 30% B; 75-80 min: 30 to 70% B; 80-85 min: 70% to 100% B; 85-90 min: 100% B (solvent A: 0.1% *v*/*v* formic acid; solvent B: 0.1% formic acid in 80% *v*/*v* acetonitrile). Mass spectra were acquired on a Thermo Scientific Q-Exactive HF Orbitrap mass spectrometer with the following settings: polarity, positive; data independent acquisition (DIA); MS resolution, 120,000; maximum ion injection time, 230 ms; MS automatic gain control (AGC) target, 3e6; normalized collision energy (NCE), 30; MS2 resolution, 30,000; MS2 maximum ion injection time, 45 ms; isolation window, 11 m/z; and MS2 AGC target, 1e6. DIA mass spectrometry data was searched using MSFragger v.3.8 using typical DIA settings against UniProt Swiss-Prot database ^77^ retrieved using Philosopher v.4.8.1 ^78^ on 2023-06-27 with added contaminants. The search results were post-processed using MSBooster v.1.1.6 followed by Percolator v.3.

Peptide identification was accepted at 1% Percolator FDR. Mass isotopomer intensities were quantified using Riana and protein turnover rate constants calculated as described above.

### Human primary fibroblast protein turnover kinetics and secretome analysis

All experiments utilizing human tissue were performed in accordance with COMIRB Protocol #01-568 at the University of Colorado Anschutz Medical Campus. As previously described ^85^, primary human CFs were isolated from left ventricular tissue obtained from non-failing donor hearts not suitable for transplant by enzymatic digestion (100 mg collagenase type 2 [Worthington Biochemical Corporation]; 1 mg trypsin [Worthington TRL3]; and 15 mg bovine serum albumin [Sigma A5611] reconstituted in 40 mL of DMEM. Fibroblasts were maintained in FGM™-3 Cardiac Fibroblast Growth Medium 3 BulletKit™ media (Lonza CC4526) and cultured to Passage 3. Fibroblasts were plated in 6-well plates at 80,000 cells/well and cultured for 2 days in FGM™-3 medium, and subsequently cultured for 48 hours in low serum Fibroblast Basal Medium (CC-3131) containing 6% D_2_O. This conditioned media was collected and processed with Seer Proteograph XT as above. Cleaned peptides from both NP mixtures were reconstituted in 0.1% formic acid to a final concentration of 0.1025 µg/µL. Resuspension concentrations were selected based on available mass of the digested peptides. In parallel, the intracellular samples were harvested and digested as described above. The samples were then analyzed using DIA mass spectrometry as above.

### Additional data analysis

Subcellular localizations were predicted using DeepLoc 2.0 ^86^ or UniProt ^77^ annotations. Over-representation and gene set enrichment analyses were performed using the ReactomePA ^87^ and fgsea packages in R and using StringDB ^88^. Skewness of protein turnover rate distributions of secreted protein is calculated as Pearson’s moment coefficient of skewness, and the bootstrap significance is calculated as the proportion of samples with more skewed distributions among 1e6 random samples with replacement of the entire data set, with sample sizes matched to the annotations being compared. Degron motifs are retrieved from Degronopedia (2024-02-01 release) ^46^. Protein energetic cost is calculated using the ECPA method with human/heterotroph H11 cost values ^45^. Protein copy number per cell is calculated using the proteomic ruler ^89^ method from the MS1 label free quantity of peptides in labeled and unlabeled samples, integrated from m_0_ to m_5_ using Riana:

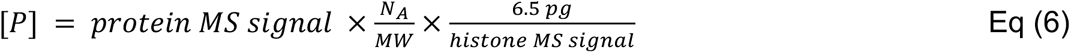

The predicted protein copy numbers are compared with prior per-protein values in mouse embryonic fibroblasts ^12^ and histone proportion per cell ^89^. The total protein mass and cell mass predicted from the proteomic ruler is also in range with values predicted from geometry. Briefly, we assume hiPSC to be spherical cells between 10–12 µm in diameter. Assuming a cellular density of 1.06 g/cm^3^ yields cell mass of 0.555–0.959 ng. Dry mass is assumed to be 30% of cellular mass and protein mass 60% of total dry mass based on literature estimates ^90^.

## Data and Code Availability

Riana is open-source and freely available on GitHub at http://github.com/ed-lau/riana. A visualization web app for isotopomer envelopment is available at http://heart.shinyapps.io/D2O_Isotope/. Raw mass spectrometry data will be available on ProteomeXchange.

## Supplemental Data

**Table S1.** Predicted average number of deuterium exchangeable hydrogen sites in each cell line, alongside commonly used adult animal values.

**Table S2.** Relative intensity of labeled peaks of selected amino acids following acid hydrolysis.

**Table S3.** Protein turnover rates from control AC16 cells in single-time-point D_2_O labeling experiments.

**Table S4.** Protein turnover rates from hiPSC-CM in single-time-point D_2_O labeling experiments.

**Table S5.** Protein turnover rates from hCF in single-time-point D_2_O labeling experiments.

**Table S6.** Protein turnover rates from thapsigargin-treated AC16 cells in single-time-point D_2_O labeling experiments.

**Table S7.** Protein turnover rates from hiPSC in time-course D_2_O labeling experiments.

**Table S8.** Gene set enrichment analysis results of fast (top 5 percentile) turnover proteins in hiPSC.

**Table S9.** Gene set enrichment analysis results of moderate (5–25 percentile) turnover proteins in hiPSC.

**Table S10.** Gene set enrichment analysis results of slow (bottom 75 percentile) turnover proteins in hiPSC.

**Table S11.** Protein turnover rates from hiPSC upon early stages (0–24 hours) of pluripotency exit.

**Table S12.** List of secreted proteins with high secretory flux that are in hiPSC-CM, hCF, or both cell types.

## Supplemental Figures

**Figure S1.**
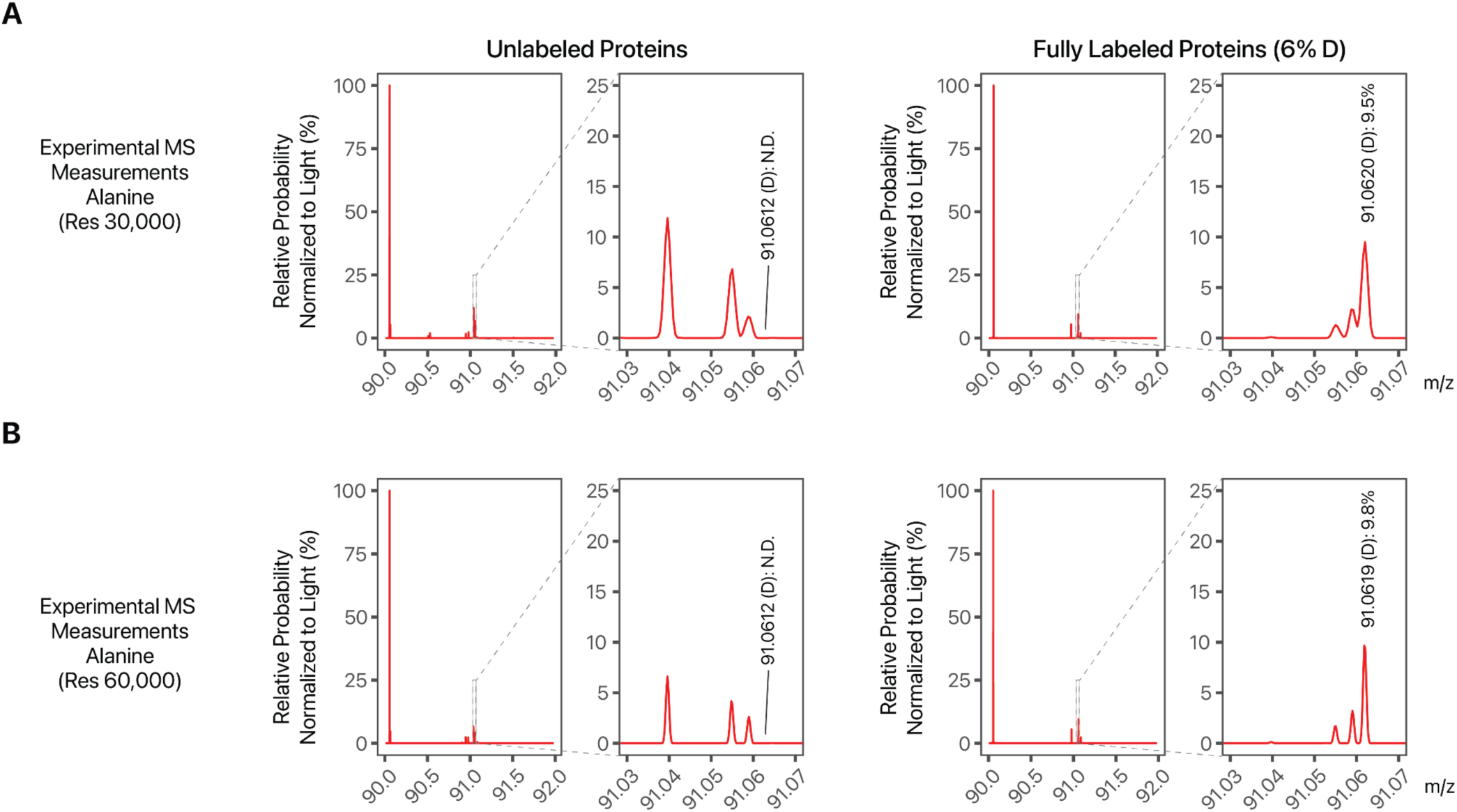
Amino acid analysis results at different resolution. **A.** MS1 mass spectra of acid hydrolysis-released amino acids from unlabeled and fully labeled AC16 cells at 30,000 resolution. Spectra are summed over 1 minute of direct infusion (see Methods). Experimental mass spectrometry data from acid hydrolysis of proteins from AC16 cells that are unlabeled (left) and fully labeled (right) with 6% D_2_O, measured at 30,000, as compared to 120,000 resolution in the main text. Upon 6% D_2_O, the relative intensity of alanine containing one D (m/z 91.0612) increased to 9.5%, consistent with 1 – 2 labeling sites in cell culture as opposed to 4 labeling sites in adult mice in vivo. The ^13^C and D containing isotopomers of alanine were not fully separable in 30,000 resolution. **B.** Same as A, but at 60,000 resolution; D-alanine relative intensity is measured at 9.8%, consistent with predicted labeling in vitro.

**Figure S2.**
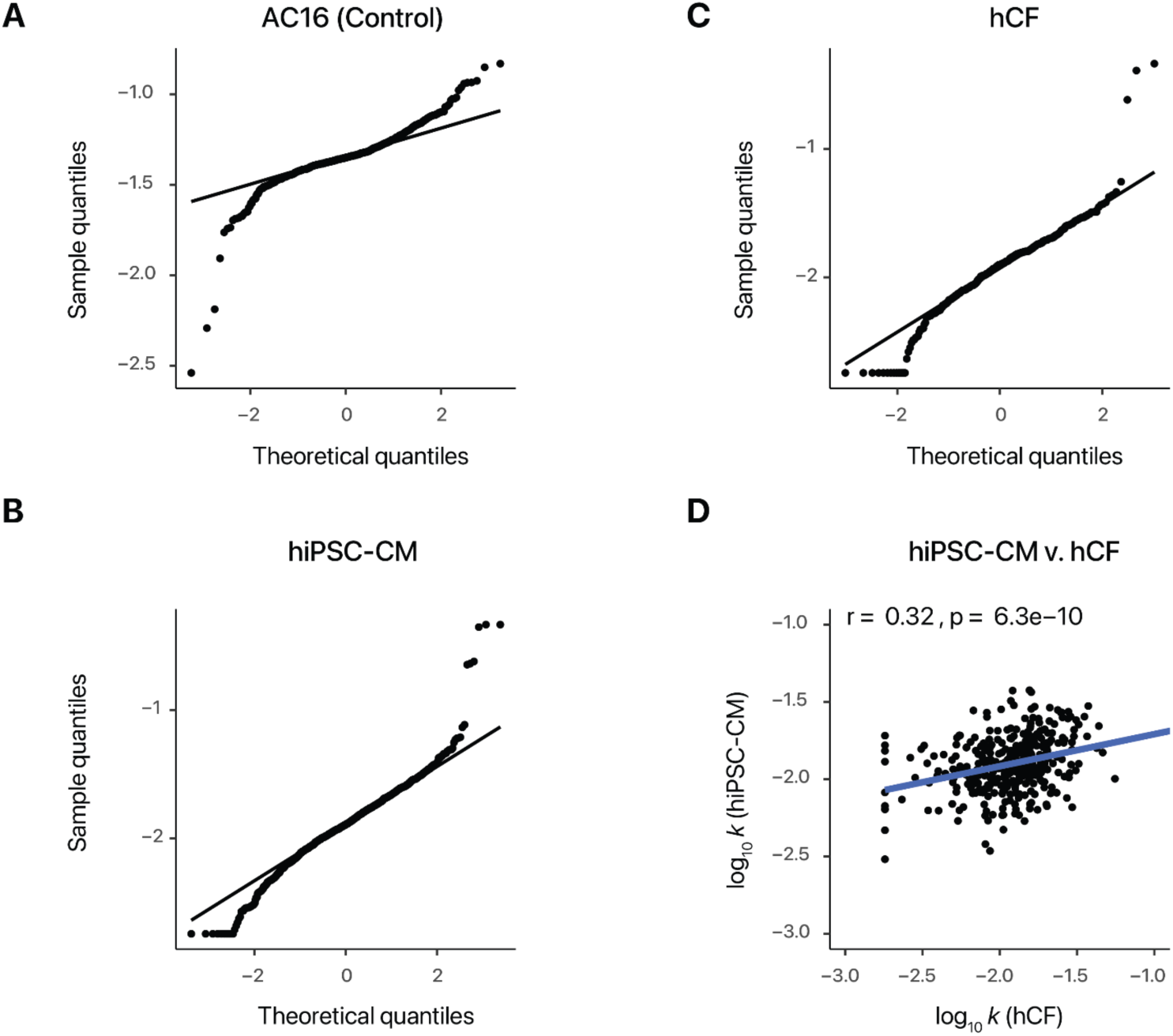
Distribution of protein turnover rates in single-time-point experiments. **A.** Quantile-quantile plots of log_10_ protein turnover rates in AC16 cells measured from D_2_O labeling **B.** Quantile-quantile plots of log_10_ protein turnover rates in hiPSC-CM cells measured from 48 hours of D_2_O labeling **C.** Quantile-quantile plots of log_10_ protein turnover rates in primary hCF cells measured from 48 hours of D_2_O labeling **D.** Moderate correlation between log_10_ turnover rates in hiPSC-CM and hCF (r: 0.32; P: 6.3e–10).

**Figure S3.**
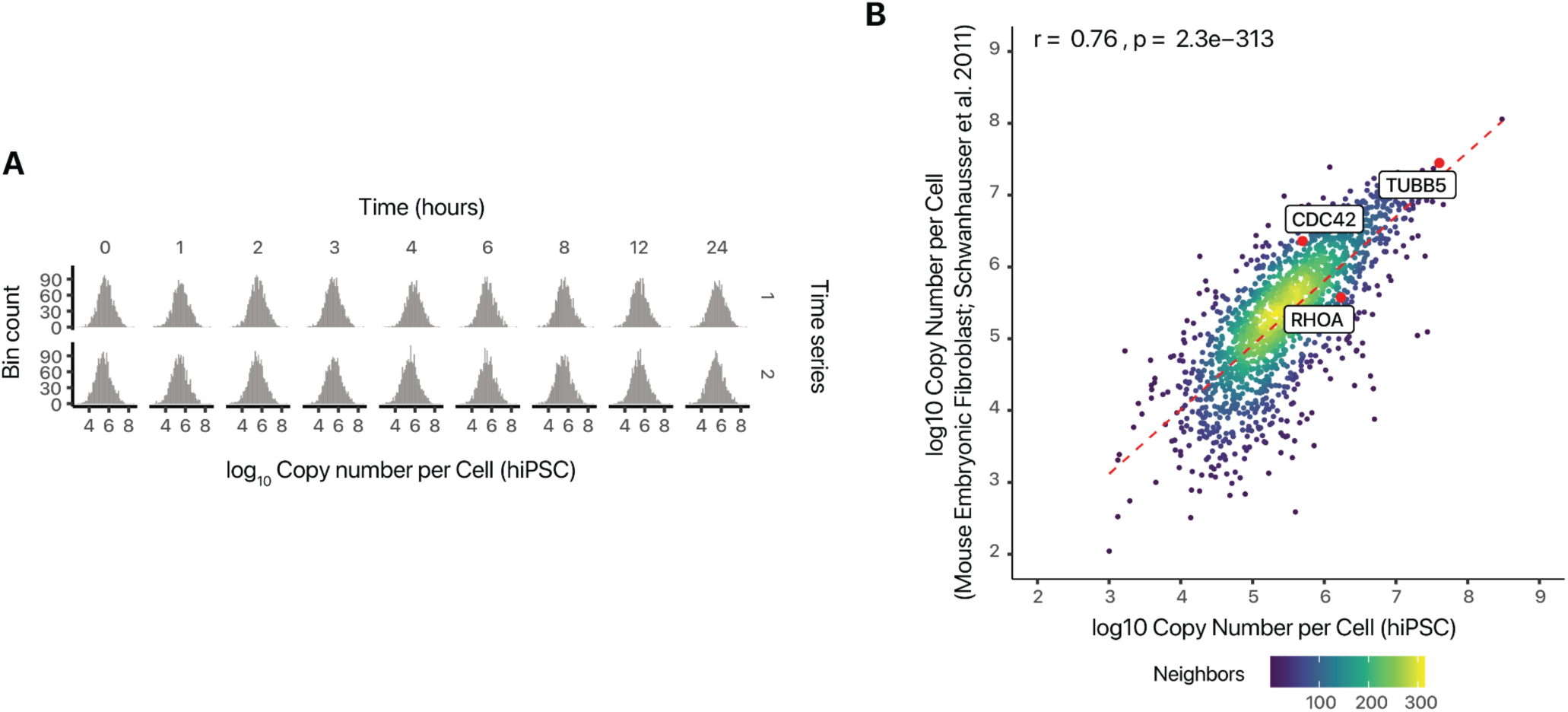
Estimation of absolute protein copy number. **A.** Protein copy number is estimated using the proteomic ruler method (Methods). Histograms show consistent distribution of protein copy number across labeling time points and time series. **B.** Scatterplot showing consistent estimates of protein copy numbers in hiPSC with prior estimates in mouse embryonic fibroblasts over nearly 5 orders of magnitude. Proteins TUBB5, CDC42, and RHOA, whose absolute abundance was further corroborated by alternative methods in Schwanhausser et al. 2011, are highlighted. Pearson’s correlation coefficient r and p value in log-log scale are shown.

**Figure S4.**
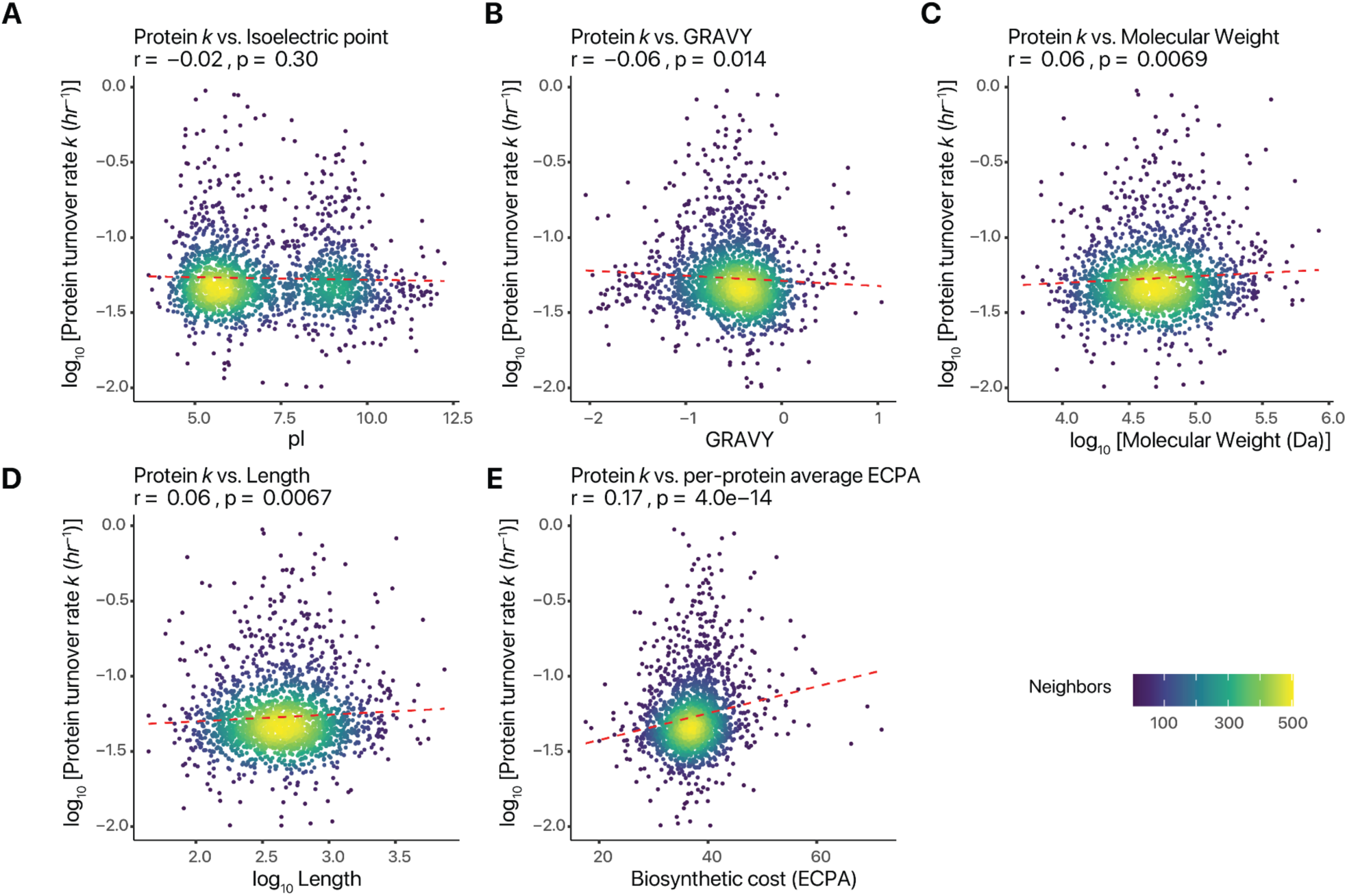
Relationship between protein turnover and sequence parameters. **A.** Scatterplot of log_10_ protein turnover rate against isoelectric point (pI). Red dashed line shows the best linear fit. Pearson’s correlation coefficient and p value in log-linear scale are shown. Colors denote point density. **B.** Same as A, but for protein hydropathy index (GRAVY). **C.** Same as A, but for protein molecular weight (Da). Pearson’s correlation coefficient and p value in log-log scale are shown. **D.** Same as C, but for protein sequence length (aa) in log10 scale. **E.** Same as A, but showing a significant positive relationship with average amino acid biosynthetic energy cost (ECPA; Zhang et al. 2018).

**Figure S5.**
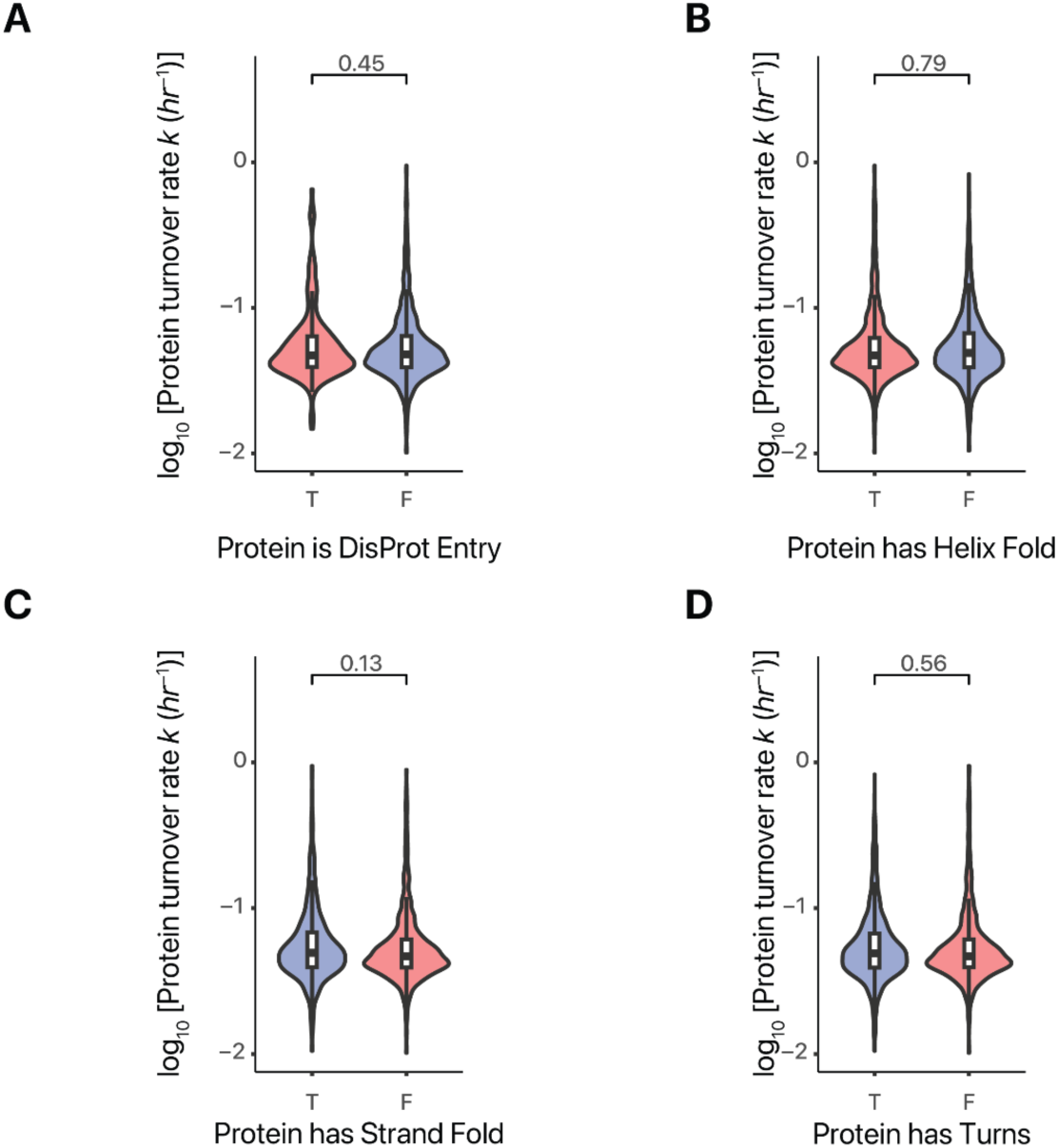
Relationship between protein turnover and gross structural feature. **A.** Violin/boxplot (IQR/1.5 IQR) showing the distribution of log10 turnover rates among proteins found in DisProt and those not found in DisProt. P value of two-tailed unpaired t-tests is shown. **B.** Same as A, but for whether a protein’s structure contains one or more annotated helix folds in Uniprot. **C.** Same as B, but for whether a protein’s structure contains one or more annotated strands in UniProt **D.** Same as C, but for whether a protein’s structure contains annotated turns in UniProt.

**Figure S6.**
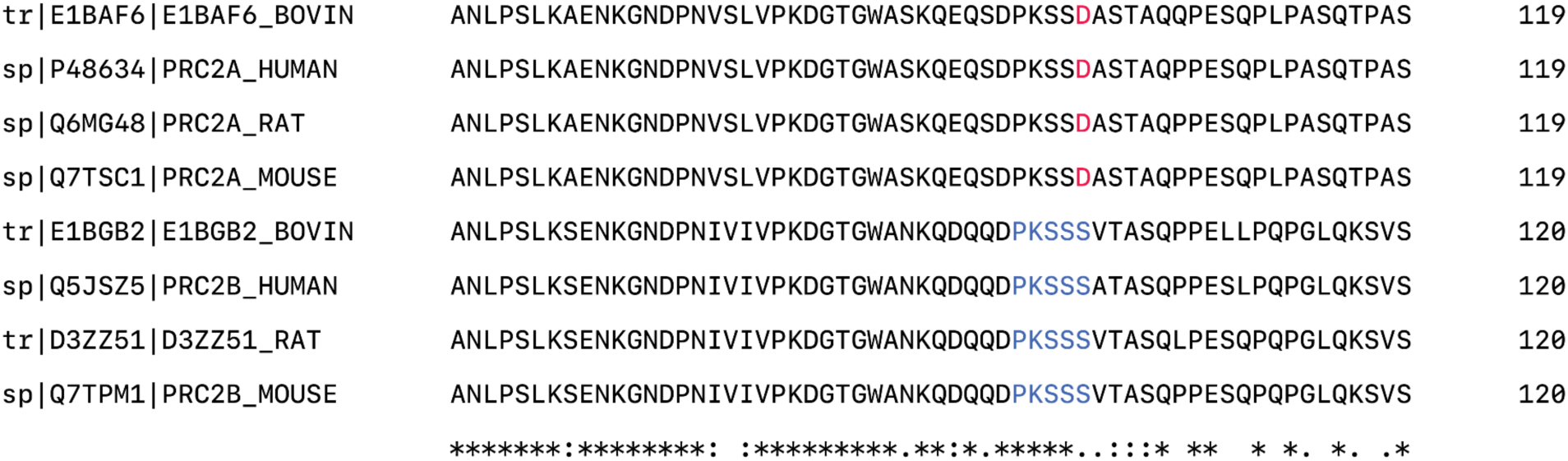
Evolutionary conservation of the PRRC2B N-terminus degron. The UniProt accession and aligned sequences surrounding the PRRC2B N-terminal SPOP degron are shown. Blue: SPOP motif in PRRC2B.

**Figure S7.**
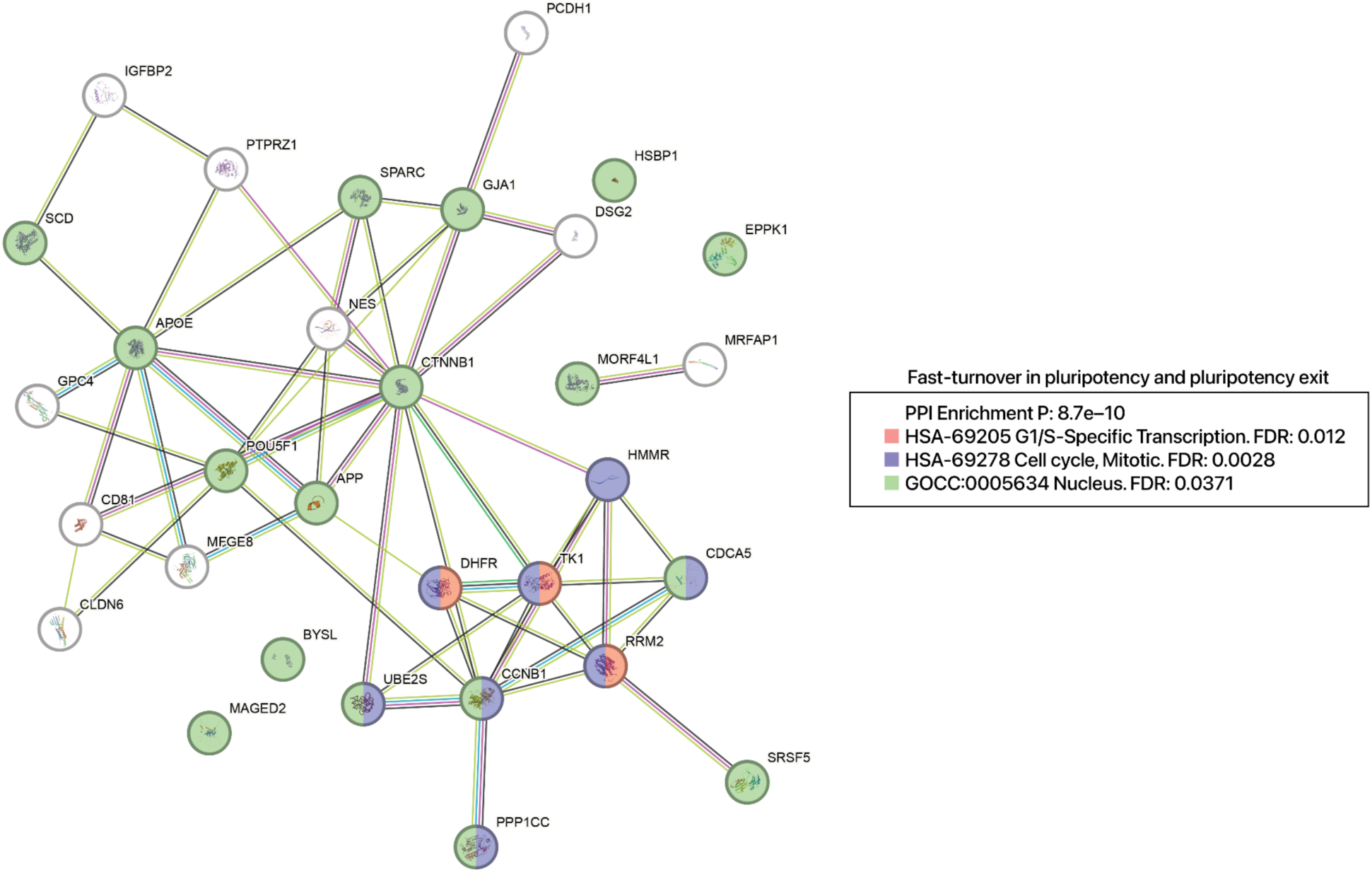
Hyperdynamic hiPSC proteins upon pluripotency exit. STRING graph showing a module of nuclear and cell cycle related proteins, which have maintained fast turnover rates (i.e., top 10 percentile of *k*) in both pluripotent and differentiating hiPSC. Colors highlight several significantly enriched Reactome and Gene Ontology terms over the genomic background.

**Figure S8.**
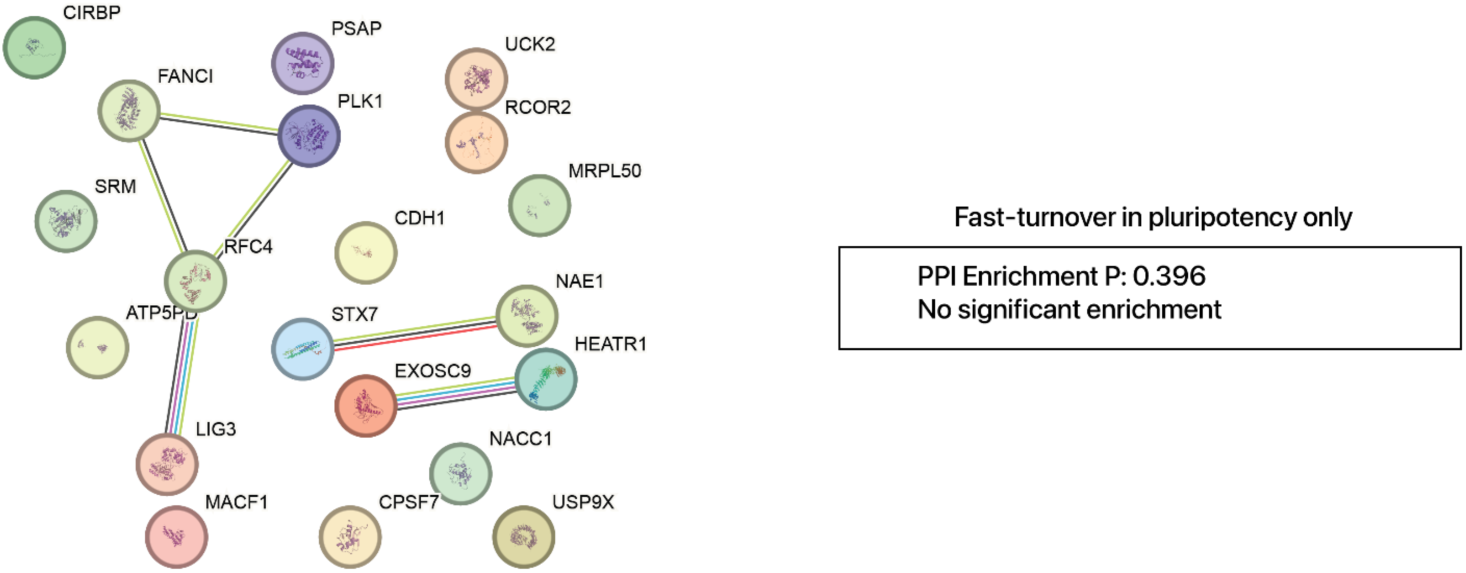
Gain of new fast-turnover proteins during early mesendoderm induction As in FIgure S7, but for proteins that are fast-turnover in pluripotent cells but become presumably parsimoniously synthesized upon pluripotency exit.

**Figure S9.**
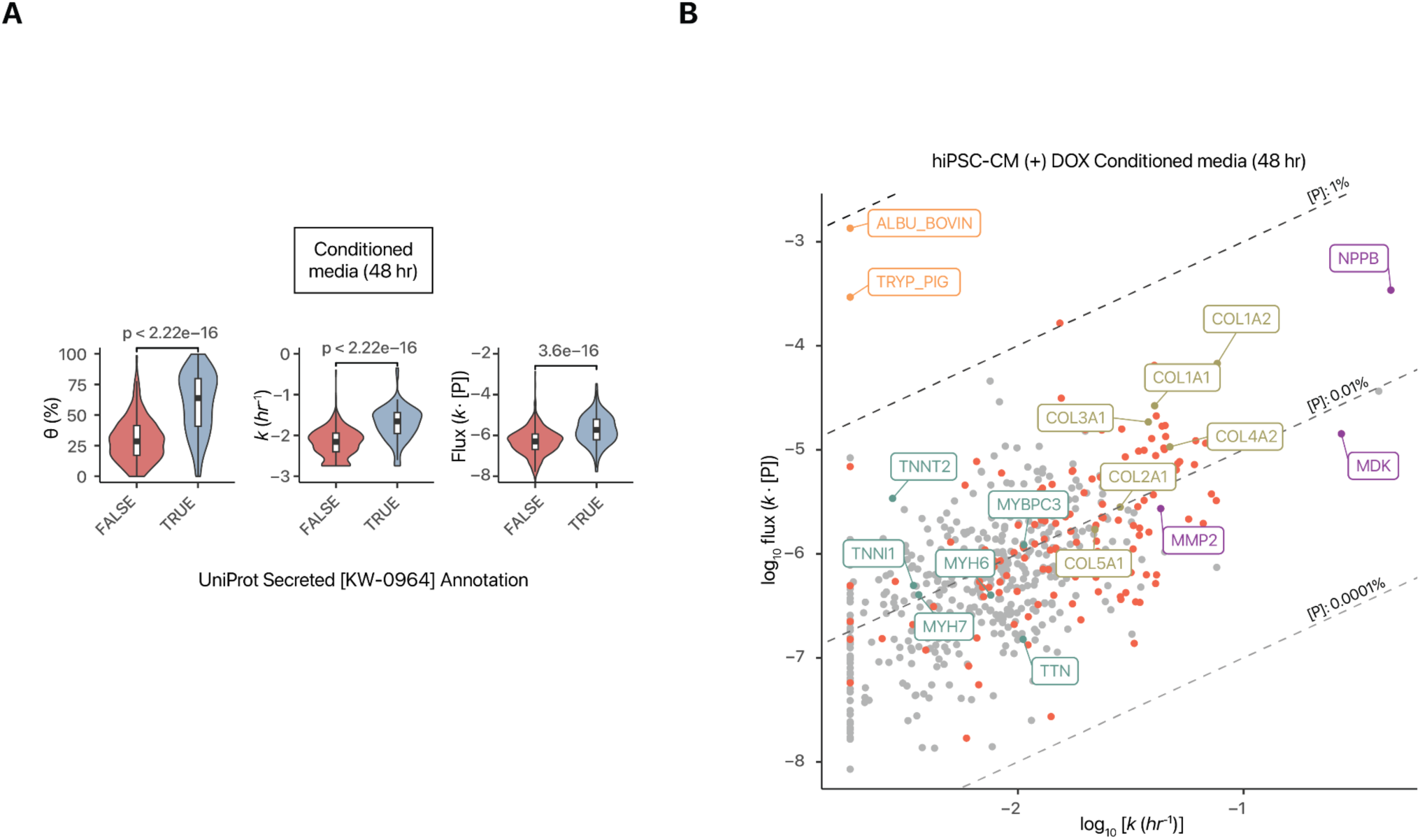
Secretome kinetics in doxorubicin-treated hiPSC-CM. **A.** Violin/boxplots (IQR/1.5-IQR) showing the fractional synthesis (left); *k* (center) and flux (right) of proteins in the conditioned media of DOX-treated hiPSC-CM after 48 hours of labeling, corresponding to the DMSO data in Figure 6C. **B.** Scatterplot showing the relationship between turnover rates and flux of proteins identified in the conditioned media of DOX-treated hiPSC-CM at 48 hours of labeling. UniProt Secreted proteins are in red. Colored labels refer to groups of highlighted proteins as in Figure 6E.

